# New insights into the 17β-hydroxysteroid dehydrogenase type 10 and amyloid-β 42 derived cytotoxicity relevant to Alzheimer’s disease

**DOI:** 10.1101/2025.02.21.639531

**Authors:** Aneta Houfková, Monika Schmidt, Ondřej Benek, Ivo Fabrik, Rudolf Andrýs, Lucie Zemanová, Ondřej Soukup, Kamil Musílek

**Affiliations:** University of Hradec Kralove, Faculty of Science, Department of Chemistry, Rokitanskeho 62, 500 03 Hradec Kralove, Czech Republic; University Hospital Hradec Kralove, Biomedical Research Centre, Sokolska 581, 500 03 Hradec Kralove, Czech Republic

**Keywords:** 17β-hydroxysteroid dehydrogenase type 10, amyloid-β peptide, Alzheimeŕs disease, mitochondria, enzyme inhibition

## Abstract

The multifunctional mitochondrial enzyme 17β-hydroxysteroid dehydrogenase type 10 (HSD10) plays an important role in the pathology of several diseases, of which Alzheimer’s disease (AD) is the most debated. HSD10 overexpression and its interplay with amyloid-β peptide (Aβ) are considered a factor contributing to mitochondrial damage and neuronal stress observed in AD patients. This study confirms that individual overexpression of HSD10 or APP (amyloid precursor protein that gives rise to Aβ) leads to cytotoxicity, and both pathological conditions are linked to mitochondrial damage. However, the metabolic changes caused by these two overexpressions significantly differ, particularly in their effect on the tricarboxylic acid cycle and β-oxidation. Furthermore, the enzymatic activity of HSD10 is identified as the primary factor of HSD10 cytotoxicity, which is significantly exacerbated in an Aβ-rich environment and can be partially reversed by HSD10 inhibitors. Notably, a previously published and competitive benzothiazole inhibitor was effective in restoring the viability of HSD10 overexpressing cells alone and in an Aβ-rich environment, implying the potential benefit of HSD10 inhibitors in mitochondrial diseases and/or AD treatment.

**GRAPHICAL ABSTRACT:** 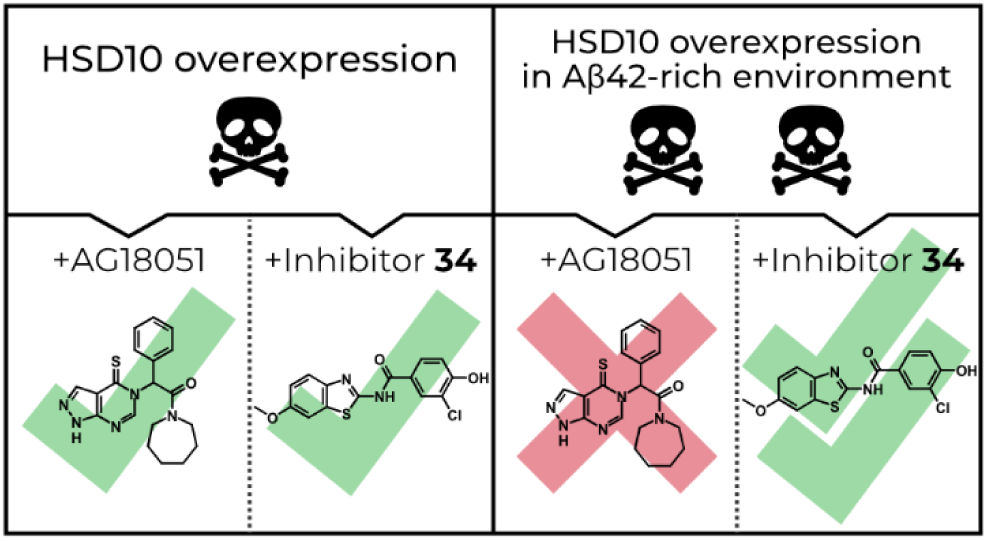

## INTRODUCTION

17β-hydroxysteroid dehydrogenase type 10 (HSD10, UniProt: Q99714), a member of the short-chain dehydrogenase/reductase superfamily, is a multifunctional mitochondrial NAD^+^-dependent enzyme involved in fatty acid metabolism, isoleucine degradation, and steroid metabolism.^1^ Besides its enzymatic activity, HSD10 is also involved in mitochondrial tRNA maturation and its further downstream processing.^2–4^ HSD10 protein was found essential for normal mitochondrial function and neuronal development, as mutations in the HSD10 gene impacting the proteińs structural stability have been associated with a rare X chromosome-linked inborn error of metabolism which causes rapid and progressive neurodegeneration prevalent in neonatal and infant age.^5–7^ The changes in HSD10 levels and/or its enzymatic function are connected with several diseases, particularly with neurodegenerative disorders such as Alzheimer’s disease (AD)^1^ and Parkinson’s disease.^8^ In AD, elevated levels of HSD10 were found in affected brain regions, where HSD10 is considered to mediate amyloid-β peptide (Aβ)-associated cytotoxicity, and this interplay is thought to be one of the possible aspects of AD pathogenesis.^9,10^

In AD progression, the engagement of HSD10 and its interplay with Aβ results in mitochondrial dysfunction and neuronal destruction.^9–11^ Aβ peptide is the cleavage product of amyloid precursor protein (APP) derived via APP subsequent amyloidogenic splicing by β- and γ-secretases. Aβ and its deposits are one of the histopathological hallmarks found in AD patients’ brains. Aβ produced by APP cleavage can range from 36 to 43 amino acids.^12^ In relation to human AD brain deposits, the Aβ42 long fragment dominates and its oligomeric forms are supposed as neurotoxic Aβ species.^13,14^

The mitochondrial dysfunction, as a consequence of HSD10 and Aβ peptide interplay, is characterized by elevated oxidative stress, increased reactive oxygen species (ROS) production, inhibition of mitochondrial complex IV, altered ATP level, the release of lactate dehydrogenase and cytochrome c from the mitochondria, DNA fragmentation, and increased apoptosis, and results in cell death.^9–11,15–17^ According to the available publications, HSD10 is thought to mediate Aβ-driven cytotoxicity, probably in a manner where Aβ affects HSD10 enzymatic function.^16–18^ The HSD10-Aβ interaction was studied using recombinant HSD10 enzyme resulting in the determination of dissociation constant 55.8 nM for Aβ42.^10^ Based on the literature, the prevention of the HSD10-Aβ interplay and/or modulation of HSD10 enzymatic function offers potential opportunities for drug development.^19–21^ To date, several classes of HSD10 inhibitors or inhibitors of HSD10-Aβ interaction have been published^19,22–26^.

However, further development and evaluation of the aforementioned HSD10 inhibitors require a better understanding of their potential to mitigate the HSD10 overexpression-related pathology. Thus, a comprehensive understanding of the HSD10 overexpression role in AD and its cellular consequences is essential for objective evaluation. This study hence aimed to describe the effects of individual HSD10 and APP overexpression on cellular fitness with the main focus on HSD10’s role in cytotoxicity development, both independently and in the presence of Aβ. In addition, the acquired findings were used to develop a novel model for testing HSD10 inhibitors, enabling the scoring of their ability to reduce pathology linked to HSD10 and Aβ, two critical factors in AD progression.

## RESULTS AND DISCUSSION

Besides its physiological functions, mitochondrial enzyme 17β-hydroxysteroid dehydrogenase type 10 (HSD10) is also connected with neurodegeneration, e.g. Alzheimer’s disease (AD), Parkinson’s disease, or HSD10 disease. In AD, HSD10 was found to exacerbate the amyloid-β peptide (Aβ) pathology, leading to enhanced cell stress and mitochondrial dysfunction.^9,16^ To better understand the role of the HSD10 enzyme in the pathology of AD development, stably-expressing monoclonal cell lines of human embryonic kidney HEK293 cells were created, overexpressing either HSD10 (referred to as HSD10 cells; Figure 1A, Supporting Information Figure S1-S2) or APP_Swe/Ind_ protein (amyloid-β precursor protein isoform 695 harbouring Swedish and Indiana mutations; referred to as APP_Swe/Ind_ cells; Figure 2A, Supporting Information Figure S4-S5). These cell lines allowed description of the consequences of individual overexpression in detail. Moreover, a catalytically inactive HSD10 mutant cell line (Y168, K172 – mutations in the catalytic active site;^18,27^ referred to as HSD10_mut_; Figure 1A, Supporting Information Figure S1-S2) was also constructed to reveal the importance of the HSD10 enzymatic activity in related processes. The loss of enzyme activity was confirmed using the recombinant HSD10 enzyme and E2 as a substrate (data not shown). Although the HEK293 cell line is not a neuronal cell line, it was chosen because the cells are easy to transfect and has many times been proposed as an AD model^28–31^ exhibiting some neuron characteristics and expression of neuronal genes. ^32,33^

**Figure 1:**
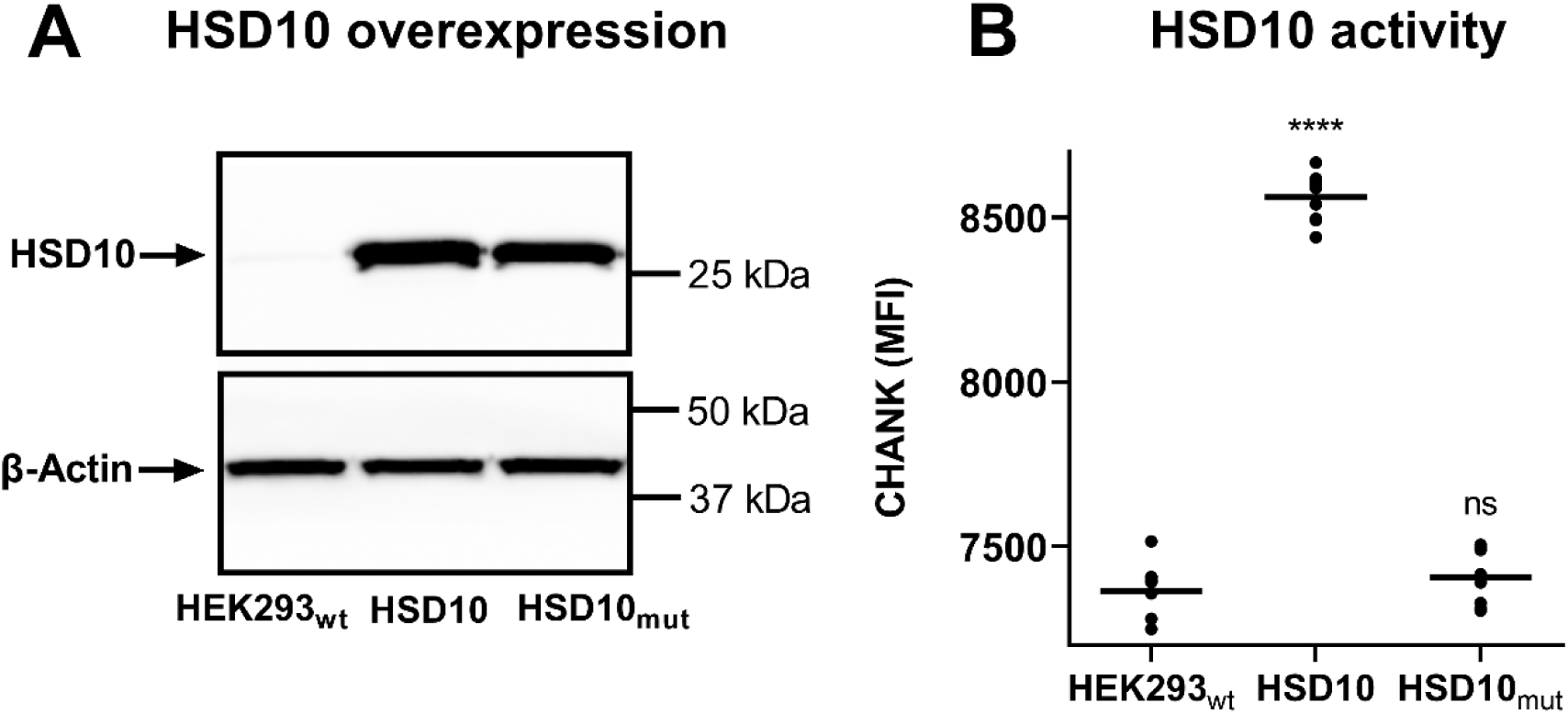
HSD10 overexpression in HSD10 and HSD10_mut_ cells. (A) Immunoblotting analysis in HEK293_wt_, HSD10, and HSD10_mut_ cells. (B) HSD10 activity determination via (-)-CHANA to CHANK turnover in HEK293wt, HSD10, and HSD10_mut_ cells (Ex/Em=380/525nm). Statistical difference: \**p* ≤ 0.05, \*\**p* ≤ 0.01, \*\*\**p* ≤ 0.001, \*\*\*\**p* ≤ 0.0001 compared to the HEK293_wt_ control group.

**Figure 2:**
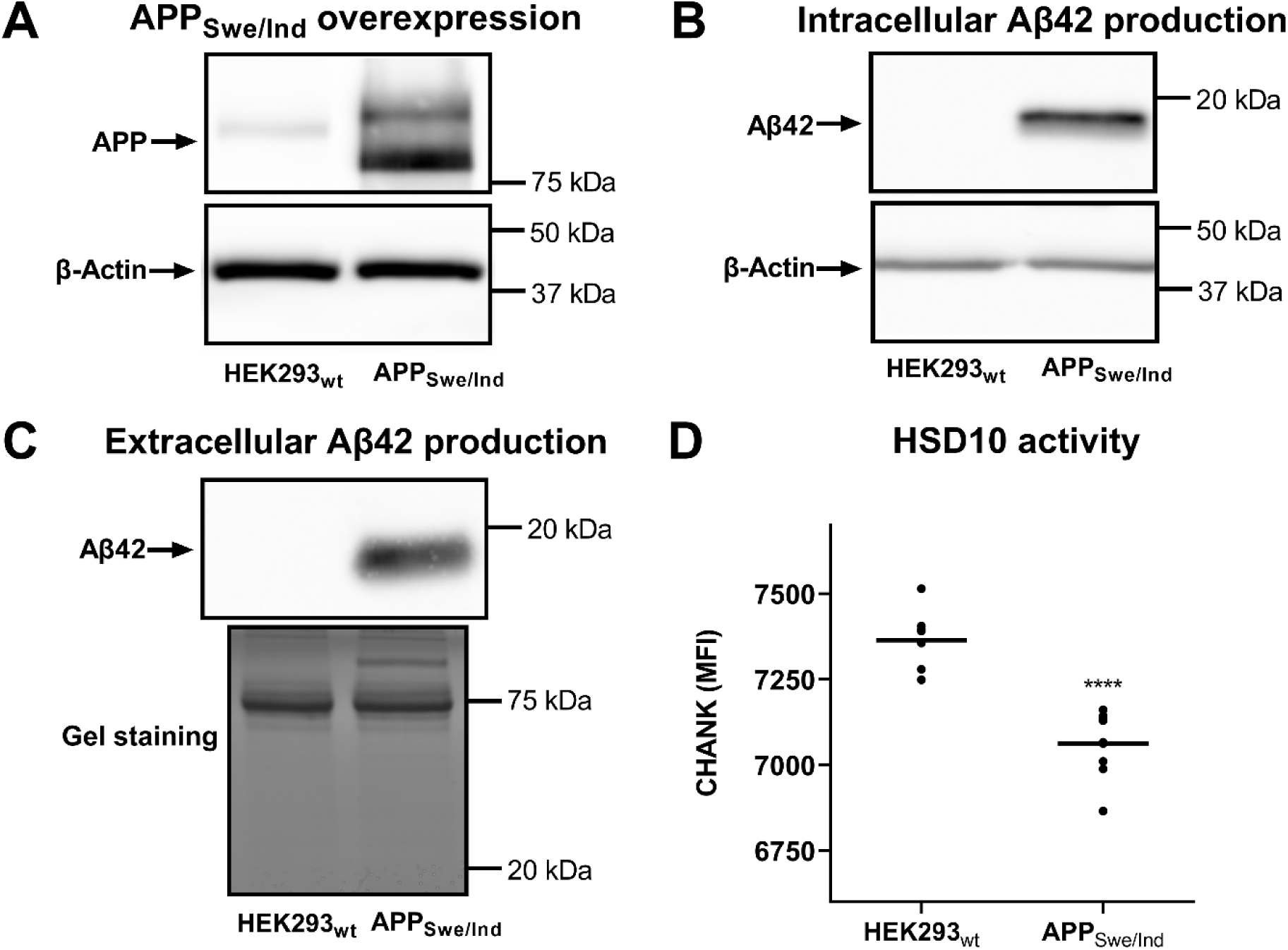
APP overexpression in APP_Swe/Ind_ cell line. (A) Immunoblotting analysis of APP expression in HEK293_wt_, and APP_Swe/Ind_ cells. (B) Immunoblotting analysis of Aβ42 production in HEK293_wt_, and APP_Swe/Ind_ cells. (C) Immunoblotting analysis of Aβ42 secretion to cultivation medium by HEK293_wt_, and APP_Swe/Ind_ cells (non-concentrated sample detection). (D) HSD10 activity determination via (-)-CHANA to CHANK turnover in HEK293_wt_, and APP_Swe/Ind_ cells (Ex/Em=380/525nm). Statistical difference: \**p* ≤ 0.05, \*\**p* ≤ 0.01, \*\*\**p* ≤ 0.001, \*\*\*\**p* ≤ 0.0001 compared to the HEK293_wt_ control group.

### HSD10 mutations in the active site abolish the enzymatic activity

HSD10 protein besides its enzymatic activity is also a structural component of the mitochondrial tRNAs processing complexes.^2,4^ To distinguish the enzymatic and non-enzymatic role of the HSD10 protein, we evaluated the HSD10 and HSD10_mut_ monoclonal cells for their ability to convert the steroid-like (-)-CHANA fluorogenic probe^34^ as a specific substrate for HSD10 enzyme in the cellular environment. The HSD10 cells demonstrated elevated fluorogenic probe conversion that implied an increase in HSD10 enzyme activity compared to the wild-type HEK293 cell line (HEK293_wt_; Figure 1). Contrastingly, the HSD10_mut_ cell line failed to convert the fluorogenic probe (Figure 1B, Supporting Information Figure S3) despite the presence of HSD10 protein levels in both HSD10 and HSD10_mut_ overexpressing cell lines (Figure 1A, Supporting Information Figure S1-S2). This confirms that mutations in the active site of HSD10 led to the loss of the catalytic activity.^10^

### APP_Swe/Ind_ overexpression leads to Aβ42 production and its release from the cells

Given the central role of Aβ in the pathogenesis of AD, the Aβ pathological hallmark was further investigated using an APP_Swe/Ind_ overexpressing cell line (Figure 2A, Supporting Information Figure S4-S5). The Aβ peptide was found to be produced inside APP_Swe/Ind_ cells (Figure 2B, Supporting Information Figure S6) and its presence was also found in APP_Swe/Ind_ cell-conditioned medium (Figure 2C, Supporting Information Figure S7-S8) which demonstrated not only the cell-production of the Aβ but also its release from the APP_Swe/Ind_ cells (in the nanomolar range). To confirm the production of the specific Aβ42 fragment (fragment production driven by the Swedish and Indiana mutations), two different antibodies for Aβ immunodetection were used. A signal around 20 kDa was observed for the antibody binding to all Aβ lengths (clone D54D2), whereas no signal was observed for the monoclonal antibody specific for Aβ40 (clone 32A1; data not shown), ruling out the possibility of Aβ40 production.

The presence of Aβ42 was confirmed using mass-spectrometry analysis by identifying Aβ42-specific peptides in the tryptic digest of gel-separated APP_Swe/Ind_ conditioned serum and protein-free media (Supporting Information Figure S9A, B and E). Moreover, the mass-spectrometric analysis unveiled acetylation of the Aβ42 peptide on the N-terminus (aspartic acid, D, Figure 3A and 3B) indicating post-cleavage processing of Aβ42 peptide and further strengthening the evidence of intracellular Aβ42 production. Besides Aβ42, the APP_Swe/Ind_ conditioned medium also contained full-length APP_Swe/Ind_ (Supporting Information Figure S9B) and APP-derived N-terminal cleavage fragments (sAPPβ) (Supporting Information Figure S9C), as indicated by the identified sequence coverage. The oligomeric state of the cell-derived extracellular Aβ42 was determined using size-exclusion chromatography followed by the immunodetection of separated fractions, revealing the most abundant Aβ42 detection under 20 kDa (between 13 to 19 kDa) (Figure 3C, Supporting Information Figure S10-S11). Assuming the molecular weight of the monomeric form is around 4.5 kDa, it is likely that the Aβ42 peptide present in the APP_Swe/Ind_ conditioned medium assembles into oligomers rather than remaining in monomeric form.

**Figure 3:**
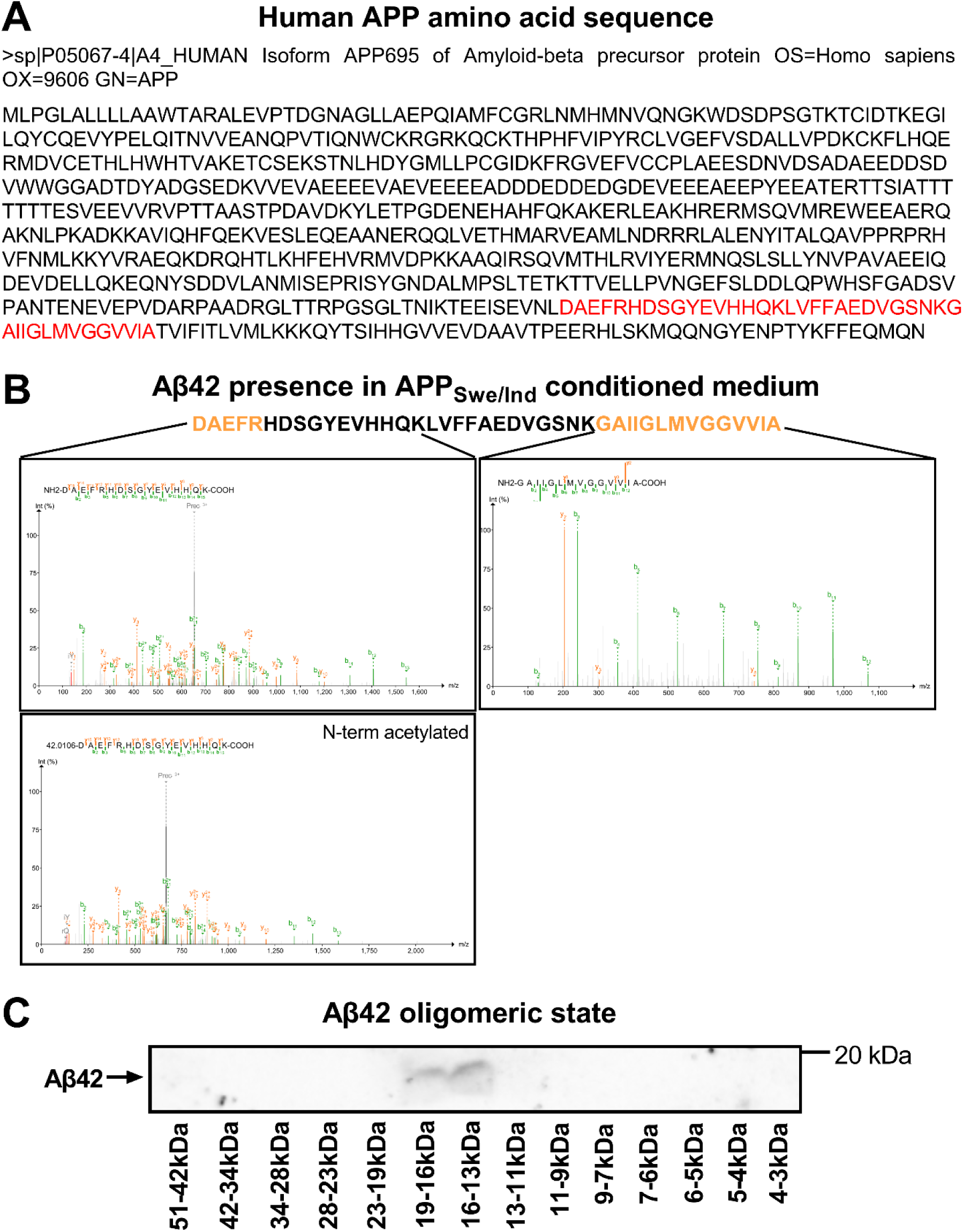
Cell-derived Aβ42 analysis. (A) The amino acid sequence of human APP_Swe/Ind_, from human reference proteome FASTA file (Uniprot) used in proteomic data search. The sequence of Aβ42 is highlighted in red. (B) Confirmation of Aβ42 presence in APP_Swe/Ind_ conditioned medium. Proteins isolated from the medium were separated by SDS-PAGE and digested by trypsin. Tryptic peptides specific for Aβ42 (yellow), and not for APP precursor, were identified only in gel bands corresponding to <20 kDa (Band E of S9A) (see representative MSMS spectra). For details see materials and methods. (C) Western blot analysis of separated fractions of APP_Swe/Ind_ cells’ conditioned medium retrieved from size-exclusion chromatography.

### APP_Swe/Ind_ overexpression affects the endogenous HSD10 enzymatic activity

The cell-based production of Aβ42 prompted an investigation into the effect of intracellular Aβ42 on HSD10 enzymatic activity. Experiments using a fluorogenic probe demonstrated that APP_Swe/Ind_ cells have reduced HSD10 enzymatic activity compared to the HEK293_wt_ cell line (Figure 2D, Supporting Information Figure S3), indicating that the APP_Swe/Ind_ overexpression associated with Aβ42 production affects the endogenous HSD10 enzymatic activity. We can speculate whether this reduction of HSD10 activity in the APP_Swe/Ind_ cells is due to Aβ42 alone or whether full-length APP or its cleavage fragments^35^ could also play a role. So far, only Aβ fragments have been shown to inhibit HSD10 enzymatic function.^10,36–39^

### Both individual HSD10 and APP_Swe/Ind_ overexpressions are highly toxic for the cells and are associated with mitochondrial damage

In AD research, the effect of HSD10 overexpression on cell fitness has been studied previously mainly in an Aβ-rich environment.^16,18,27^ However, HSD10 overexpression may also have its own mechanism affecting cell fitness and thus contribute to the AD pathology independently of Aβ. Indeed, understanding these processes could be essential for selecting the correct molecular target for drug research and development. This study aimed to evaluate the effect of individual HSD10, HSD10_mut_, and APP_Swe/Ind_ overexpression, respectively, on cellular fitness by determining associated ATP levels, cytotoxicity, and cell viability. The ATP quantification was done in complete glucose medium 72 hr and 168 hr after cell seeding and it showed significantly decreased ATP levels in HSD10 and APP_Swe/Ind_ cell lines, whereas the HSD10_mut_ cell line had unaltered ATP levels. All three cell lines displayed insignificant changes in cytotoxicity in comparison to control HEK293_wt_ cells even in the late interval (Figure 4A and 4B, Supporting Information Figure S12), and were fully viable in continuous cultivation on complete glucose media (several months, data not shown).

**Figure 4:**
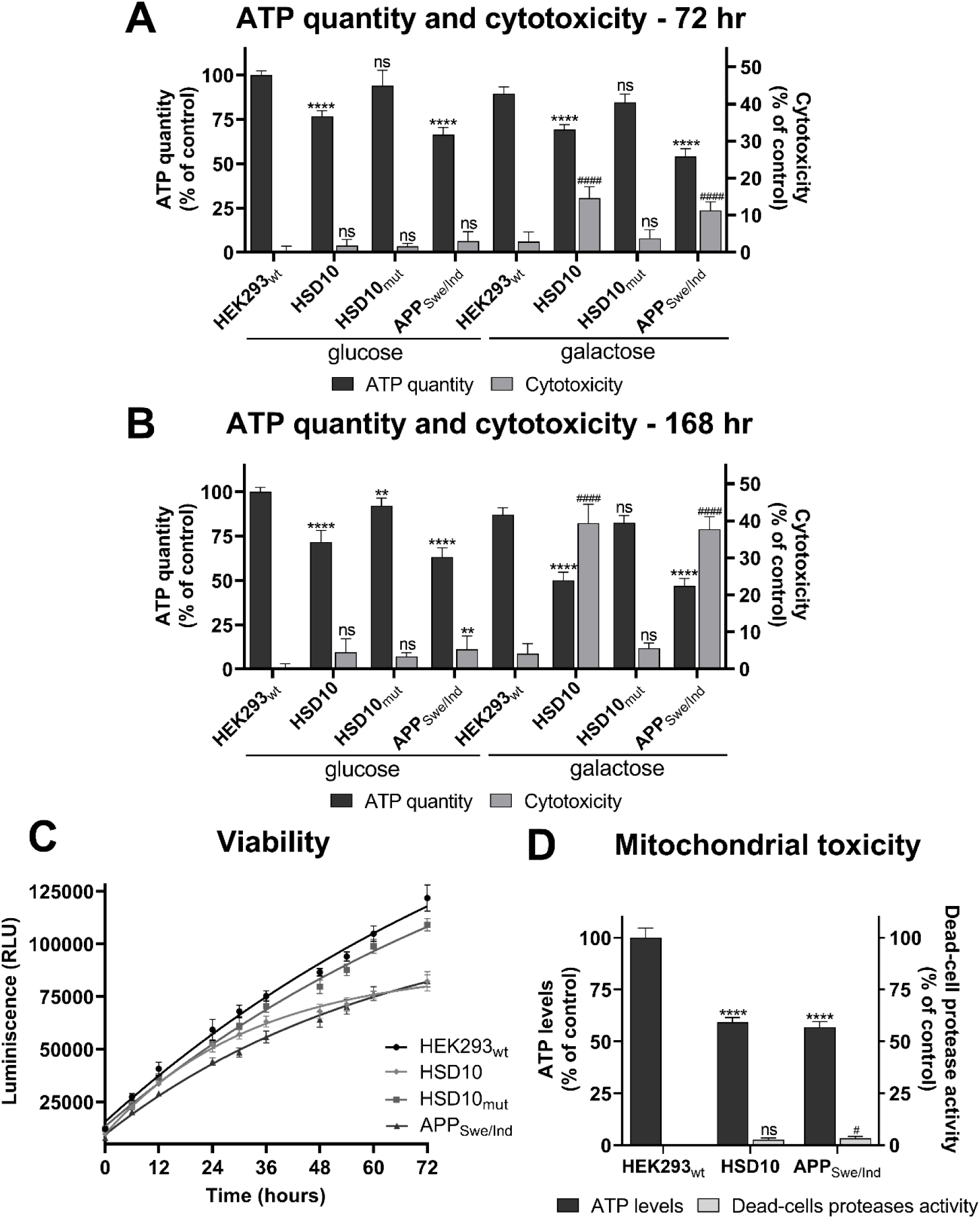
Cell and mitochondrial fitness parameters in HEK293_wt_, HSD10, HSD10_mut_, and APP_Swe/Ind_ cells. (**A**) ATP levels and cytotoxicity in HEK293_wt_, HSD10, HSD10_mut_, and APP_Swe/Ind_ cell lines 72 hr post-seeding into the glucose and galactose media. Data were normalized between DMSO-treated (1%) and valinomycin-treated (100 µM) HEK293_wt_ cells cultivated in a glucose or galactose medium. Values are given as means ± SD from three independent cell culture preparations with four technical replicates (n=12). (**B**) ATP levels and cytotoxicity in HEK293_wt_, HSD10, HSD10_mut_, and APP_Swe/Ind_ cell lines 168 hr post-seeding into the glucose and galactose media. Data were normalized between DMSO-treated (1%) and valinomycin-treated (100 µM) HEK293_wt_ cells cultivated in a glucose or galactose medium. Values are given as means ± SD from three independent cell culture preparations with four technical replicates (n=12). (**C**) Viability of HEK293_wt_, HSD10, HSD10_mut_, and APP_Swe/Ind_ cell lines monitored for 72 hr cultivation in galactose medium. Values are given as means ± SD from three independent cell culture preparations with three technical replicates (n=9). (**D**) Mitochondrial toxicity in HEK293_wt_, HSD10, and APP_Swe/Ind_ cell lines 72 hr post-seeding into galactose medium. Data were normalized between DMSO-treated (1%) and valinomycin-treated (100 µM) (for dead-cells protease activity) or antimycin A-treated (1 µM) (for ATP levels measurements) HEK293_wt_ cells cultivated in galactose medium. Values are given as means ± SD from three independent cell culture preparations with four technical replicates (n=12). SD: standard deviation. Statistical difference for ATP quantity and ATP levels: \**p* ≤ 0.05, \*\**p* ≤ 0.01, \*\*\**p* ≤ 0.001, \*\*\*\**p* ≤ 0.0001 compared to the HEK293_wt_ control group cultivated in glucose or galactose medium. Statistical difference for cytotoxicity and dead-cell protease activity: ^#^*p* ≤ 0.05, ^##^*p* ≤ 0.01, ^###^*p* ≤ 0.001, ^####^*p* ≤ 0.0001 compared to the HEK293_wt_ control group cultivated in glucose or galactose medium.

The apparent absence of cytotoxicity in both HSD10 and APP_Swe/Ind_ overexpressing cells was surprising, because Aβ peptide has been repeatedly reported to affect mitochondrial processes and to cause a decline in cell viability.^40,41^ Since both analysed overexpressions have been postulated to affect mitochondria,^16,18,41^ the potential cell impairment was examined by altering the major carbon source (galactose instead of glucose). Usually, the switch from glucose to galactose forces cellular energy production towards mitochondrial oxidative phosphorylation (OXPHOS),^42,43^ enhances mitochondrial responsiveness to toxic effects, and can help to amplify the existing mitochondrial damage. In contrast, cells cultured in glucose conditions can compensate for mitochondrial dysfunction via glycolysis and temporarily mask mitochondrial impairment. Hence glucose-free conditions may reveal mitochondrial and cellular damage that remains hidden in a glucose-rich environment with its cellular aerobic glycolytic activity.^44–46^

HSD10 and APP_Swe/Ind_ cells incubated in complete galactose media showed a further decrease in ATP levels compared to the levels found in complete glucose media. Moreover, the change in carbon source was accompanied by the manifestation of cytotoxicity as early as 72 hr, and was even more visible at 168 hr in both overexpressing cell lines (Figure 4A and 4B, Supporting Information Figure S12). This phenomenon was confirmed by real-time cell viability determination, which indicated a significant reduction in the numbers of viable cells of both HSD10 and APP_Swe/Ind_ compared to those of HEK293_wt_ cells after 72 hr of cultivation in galactose medium (Figure 4C, Supporting Information Figure S13). HSD10_mut_ cells (without HSD10 enzymatic activity) were not affected by the carbon source exchange and displayed similar growth and ATP levels as the control HEK293_wt_ cells (Figure 4A, 4B and 4C, Supporting Information Figure S12-S13). These findings suggest that HSD10 enzymatic activity is a primary factor responsible for HSD10-driven cell damage. Of note, the HEK293_wt_ cells showed no signs of cytotoxicity in the galactose environment, confirming that the HSD10 and APP_Swe/Ind_ overexpressions were the main cause of cell pathology, and ruling out the possibility of galactose toxicity.

Cytotoxicity associated with APP_Swe/Ind_ and/or Aβ42 has been published in the past.^40,41^ Moreover, there is also evidence suggesting that other APP cleavage products, such as the C-terminal 99 amino acid long fragment (C99), may contribute to cell damage through mitochondrial dysfunction.^47^ However, in our study, the C99 fragment was not detected in the APP_Swe/Ind_ sample (Supplementary Information Figure S9) suggesting its complete cleavage towards the Aβ42 peptide. The more interesting finding was the observed cytotoxicity of HSD10 overexpression alone, which has not been previously observed in studies using transgenic mice, transgenic mice cultured neurons,^16^ or primary mouse astrocytes.^27^ Previous data have pointed to HSD10 cytotoxicity only in the context of its interplay with Aβ leading to mitochondrial impairment. ^10,16,18^ However, our data demonstrates that HSD10 overexpression can cause pathological changes even in the absence of Aβ, providing new insight into HSD10-related diseases. Importantly, the induction of this phenomenon requires a glucose-free environment, which forces the cells to engage mitochondria and OXPHOS. This finding can be significant in the context of AD, as older age is associated with a decrease in glucose utilization,^48^ a phenomenon that was also observed in AD brains.^49^ Thus, our data suggest that problems in glucose utilization may serve as a triggering factor for the development of neuronal defects observed in AD.

Since the cytotoxic effects of HSD10 and APP_Swe/Ind_ overexpression were observed exclusively in the galactose environment, mitochondrial dysfunction in these cell lines was expected. To investigate this further, a mitochondrial toxicity assay was performed by measuring ATP levels (as an indicator of mitochondrial damage) and the activity of dead-cell proteases (as an indicator of secondary toxicity, such as necrosis) in a glucose-free medium. After 72 hr of cultivation in galactose medium, both HSD10 and APP_Swe/Ind_ cells showed a more than 40% decrease in ATP levels accompanied by negligible dead-cell protease activity, when compared to the HEK293_wt_ cells (Figure 4D, Supporting Information Figure S14). This suggests mitochondrial dysfunction, defined by a decrease of more than 20% in ATP levels and less than a 20% increase in dead-cell protease activity relative to control. These significant results confirmed the presence of mitochondrial damage in both HSD10 and APP_Swe/Ind_ overexpressing cell lines, leading to cytotoxic effects and decreased cell viability in a glucose-free environment. While these findings confirm the mitochondrial-based toxicity in both cell lines, the mechanisms of both overexpression-driven mitochondrial toxicities may be different and remain to be elucidated.

### HSD10 and APP_Swe/Ind_-driven cytotoxicity is linked to distinct metabolic changes

To gain more detailed insight into the metabolic differences between the two studied cell lines, their metabolism was analyzed using the MitoPlates S-1 assay, which determines the changes in mitochondrial electron flow (MEF) in the presence of particular substrates. The results uncovered the contrast in metabolic phenotypes of both HSD10 and APP_Swe/Ind_ overexpressing cells, with both cell lines showing completely different patterns of MEF changes resulting from the different substrates (Figure 5, Supporting Information Figure S15).

**Figure 5:**
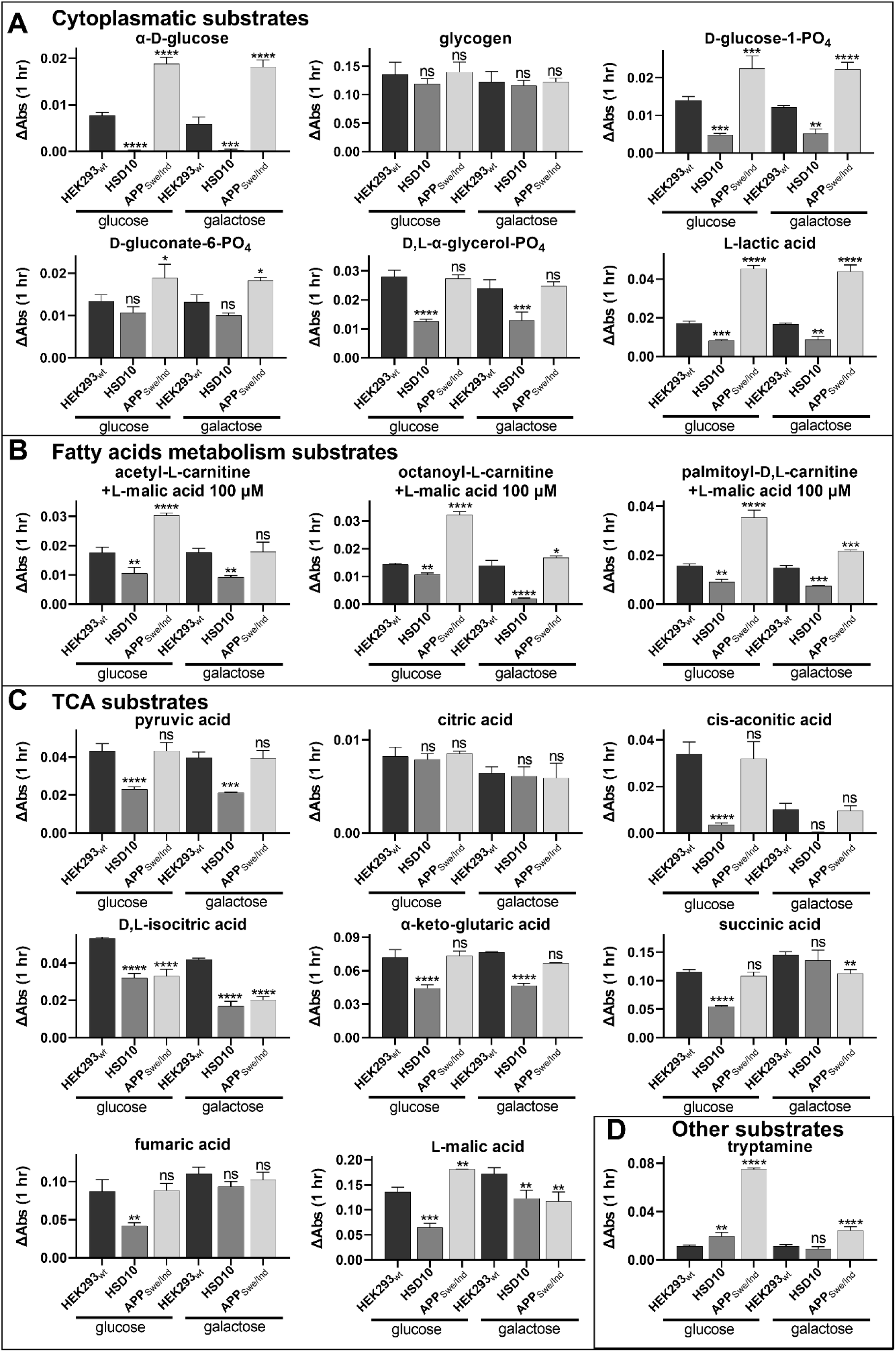
The mitochondrial electron flow changes (corresponding to metabolic changes) in HEK293_wt_, HSD10, and APP_Swe/Ind_ cells measured by absorbance (OD590) using MitoPlate S-1 assay. Conversion of cytosolic substrates (**A**), fatty acid metabolism substrates (**B**), TCA cycle substrates (**C**), and other substrates (**D**) in HEK293_wt_, HSD10, and APP_Swe/Ind_ cells. Values are given as means ± SD from three independent cell culture preparations with one technical replicate (n=3). SD: standard deviation. Statistical difference: \**p* ≤ 0.05, \*\**p* ≤ 0.01, \*\*\**p* ≤ 0.001, \*\*\*\**p* ≤ 0.0001 compared to the HEK293_wt_ control group.

HSD10 cells demonstrated a marked reduction in the MEF in the presence of α-D-glucose, D-glucose-1-PO_4_, D-gluconate-6-PO_4_, D,L-α-glycerol-PO_4_, and L-lactic acid (Figure 5A), as well as pyruvic acid (Figure 5C), which may suggest a general glucose metabolism impairment driven by HSD10 overexpression. In contrast, the APP_Swe/Ind_ cells exhibited an increase in the MEF in the presence of cytosolic substrates, including α-D-glucose, D-glucose-1-PO_4_, D-gluconate-6-PO_4_, and L-lactic acid (Figure 6A), which may indicate an overall increase in glucose metabolism due to APP_Swe/Ind_ overexpression.

**Figure 6:**
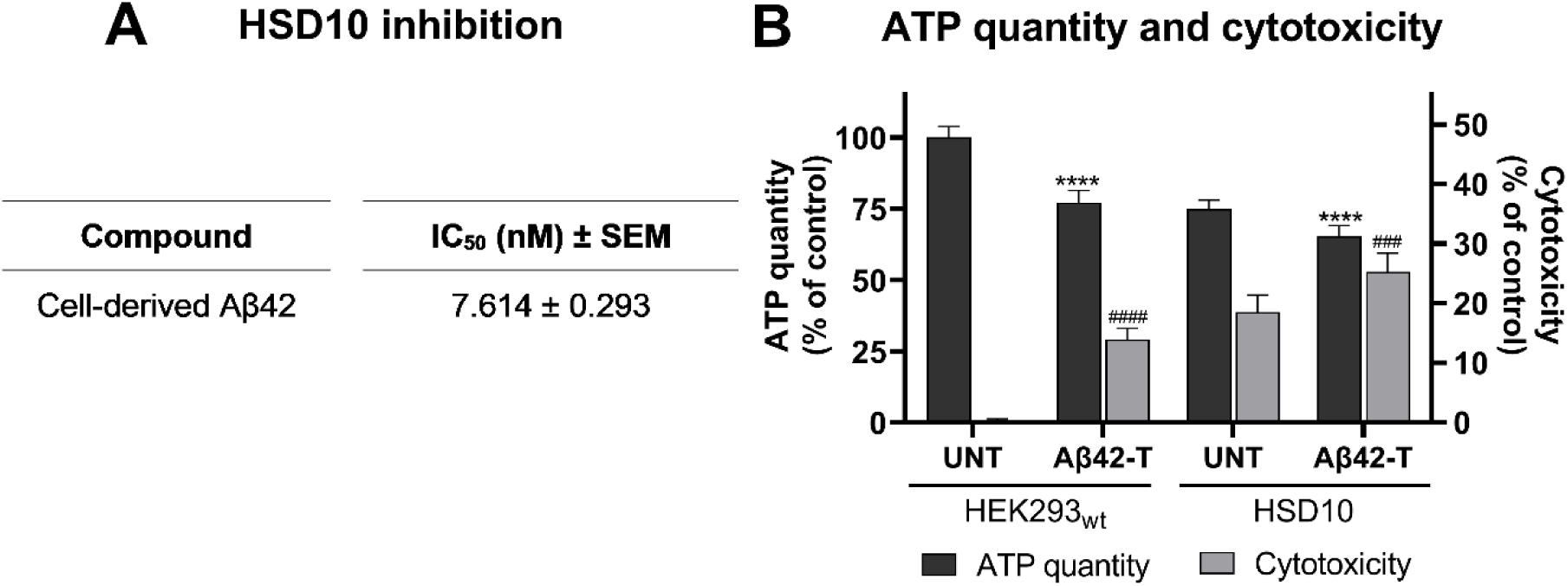
The effect of cell-derived Aβ42 treatment. (**A**) HSD10 inhibition in the cellular environment. The IC_50_ determination of cell-derived Aβ42 using (-)-CHANA fluorogenic probe in HSD10 cells in glucose medium. Values are given as means ± SEM from three cell culture preparations with three technical replicates (n=9). (**B**) ATP levels and cytotoxicity in HEK293_wt_, and HSD10 cells performed 72 hr post-seeding into galactose medium containing 7.6 nM cell-derived Aβ42. Data were normalized between DMSO-treated (1%) and valinomycin-treated (100 µM) HEK293_wt_ cells cultivated in galactose medium. Values are given as means ± SD from three independent cell culture preparations with three technical replicates (n=9). SD: standard deviation. Statistical difference for ATP quantity: \**p* ≤ 0.05, \*\**p* ≤ 0.01, \*\*\**p* ≤ 0.001, \*\*\*\**p* ≤ 0.0001 compared to the untreated HEK293_wt_ or HSD10 control group. Statistical difference for cytotoxicity: ^#^*p* ≤ 0.05, ^##^*p* ≤ 0.01, ^###^*p* ≤ 0.001, ^####^*p* ≤ 0.0001 compared to the untreated HEK293_wt_ or HSD10 control group.

Opposite patterns between the two cell lines were also observed during the processing of fatty acids (a part of the β-oxidation process; Figure 5B). HSD10 cells showed decreased MEF, which may indicate reduced β-oxidation, while APP_Swe/Ind_ cells showed upregulation of MEF during the utilization of β-oxidation substrates.

The HSD10 cells also showed reduced MEF in the presence of nearly all tricarboxylic acid (TCA) cycle substrates compared to HEK293_wt_ (Figure 5C). These results suggest a serious and overall impairment of the TCA cycle. In contrast, APP_Swe/Ind_ cells exhibited variable and substrate-specific changes (Figure 5C), indicating a less comprehensive effect of overexpression on the TCA cycle.

Both cell lines demonstrated increased MEF during tryptamine processing (Figure 5D), particularly after cultivation in a glucose medium, suggesting a probable increase in monoamine oxidase (MAO) activity. However, this increase was more pronounced in APP_Swe/Ind_ cells. Of note, the elevated MAO activity is known to be connected with enhanced reactive oxygen species (ROS) production, a phenomenon observed in AD^50,51^, and is linked to neuroinflammation, oxidative stress, and cognitive decline.^52^

It is plausible that the observed pattern of metabolic changes in HSD10 cells may be a result of increased HSD10 enzymatic activity. One of the major functions of the HSD10 enzyme is its involvement in steroid metabolism, with the most important substrate 17β-estradiol (E2) being oxidized by HSD10 to estrone.^19^ E2 has been reported to affect β-oxidation, as its supplementation led to increased β-oxidation capacity^53^, treated cells exhibited increased expression of genes related to β-oxidation^54^, and defects in expression of β-oxidation genes in a knock-out mouse model were restored after E2 treatment.^55^

Regarding the increased HSD10 enzyme activity in HSD10 cells, it is reasonable to expect increased turnover of E2 to estrone, leading to an overall decreased level of E2 in cells. This reduction in E2 levels might be a major, or at least significant factor contributing to the proposed diminished β-oxidation (Figure 5B). The decrease in β-oxidation reduces also acetyl-CoA production, which serves as an input substrate for the TCA cycle,^56^ which in turn may be linked to the suggested TCA cycle downregulation in HSD10 cells (Figure 5C). Additionally, increased MEF during tryptamine utilization (Figure 5D) might be associated with increased MAO activity in HSD10 cells and indicate that these HSD10-driven defects may be linked to elevated oxidative stress. It should be considered that the MitoPlate assay, which measures changes in MEF based on NADH and FADH_2_ production, primarily reflects activity in the mitochondrial TCA cycle. Thus, the pronounced alterations in the TCA cycle in HSD10 cells could have affected the reliability of glycolytic metabolism measurements (Figure 5A). In other words, it is unclear whether these data really reflect glycolytic activity (and a substrate other than glucose is used as the main energy source), or whether the data are biased by the significant reduction of the TCA cycle in HSD10 cells (Figure 5C).

On the other hand, the measurement of MEF in APP_Swe/Ind_ overexpression suggests a partial decline in the TCA cycle (Figure 5C) together with glucose hypermetabolism (Figure 5A). This observation aligns with previous studies. While AD is typically associated with glucose hypometabolism, evidence suggests that glucose hypermetabolism may also occur in AD patients, depending on the degree of impairment.^44,57–61^ It is hypothesized that in the early stages of the disease, cells undergo a shift from OXPHOS to aerobic glycolysis for ATP production as a protective response to Aβ-driven mitochondrial damage.^44^ As a result, cells progressively increase glucose metabolism until this shift becomes unsustainable, leading to a collapse in glucose metabolism towards reduced levels in the late stages of disease progression.^44,62^ The probable increased glucose metabolism together with the impairment of the TCA cycle observed in APP_Swe/Ind_ cells, supports previous findings and suggests that changes observed in APP_Swe/Ind_ cells may represent a protective mechanism against Aβ42-driven mitochondrial dysfunction (corresponding to the early stages of the disease). Additionally, the data also indicate a possible upregulation of β-oxidation (Figure 5B) in APP_Swe/Ind_ cells, which could be also linked to this protective response, as studies demonstrated that the shift from the TCA cycle to glycolysis may also stimulate β-oxidation.^63,64^ Finally, similarly to HSD10 cells, upregulation of mitochondrial electron flow during tryptamine utilization (Figure 5D) in APP_Swe/Ind_ cells suggests increased oxidative stress.

Taken together, the MEF measurement showed that both HSD10 and APP_Swe/Ind_ cells have different metabolic profiles. The results indicated that both overexpressions probably result in a TCA cycle impairment, although the defect in HSD10 cells appears more severe. We propose that this TCA cycle decline in HSD10 cells might be linked to increased HSD10 enzymatic activity and β-oxidation impairment, likely contributing to increased oxidative stress. In contrast, APP_Swe/Ind_ cells seemed to shift from OXPHOS to a glycolytic pathway for ATP production, possibly as a protective mechanism against Aβ42/APP_Swe/Ind_-driven TCA cycle decline. This metabolic switch is likely accompanied by β-oxidation upregulation and may also contribute to elevated oxidative stress. Although these experiments were performed using embryonic kidney cells whose metabolic profile can differ from that of neuronal cells, the HEK293 cells have repeatedly been shown as an AD model ^28–31^ since they exhibit some characteristics of neurons and express more than 60 neuronal genes, including neurofilament proteins, neuroreceptors, and ion channel subunits.^32,33^ All observations in metabolic differences between these cell lines would require further studies directly investigating the activity of involved individual enzymes. Nevertheless, these results still indicate significant differences between the HSD10 and APP_Swe/Ind_ overexpression effects on metabolic pathways, suggesting that the mitochondrial damage observed in the studied cell lines may have a different basis.

### Cell-derived Aβ42 affects cellular HSD10 activity and potentiates HSD10-driven cytotoxicity

Most studies in recent decades have used synthetic preparations of Aβ peptides to uncover the detrimental effect of these peptides at the cellular level. Due to the amphiphilic structure of Aβ, the synthetic preparation suffers from certain limitations, especially during the purification steps, the peptide handling itself, and even in the strategy to distinguish the monomeric and oligomeric forms.^39^ Nearly all papers reporting HSD10 research data were using commercially available synthetic Aβ preparation with the use of different monomerizing agents.^27,65^

In the present study, the ability of cell-derived Aβ42 to reduce HSD10 enzymatic activity inside the cells was investigated. APP_Swe/Ind_ conditioned glucose medium (as a source of Aβ42) was collected after 3 to 5 days of cell cultivation, clarified by low-speed centrifugation, quantified for the Aβ42 amount, and used for the treatment of HSD10 cells. Indeed, the effective reduction of HSD10 activity was detected by fluorogenic probe, reaching IC_50_ values of 7.6 nM (Figure 6A). These results demonstrate that APP_Swe/Ind_ overexpression and extracellular Aβ42 production can be used as effective Aβ42 peptide treatment.

Furthermore, the previous data using the same fluorogenic probe showed only moderate inhibition using a micromolar concentration of synthetically prepared Aβ42.^27,34^ This difference in results is probably caused by Aβ oligomerization status as the synthetic preparation of Aβ is known to suffer oligomeric state changes along with difficulties in its determination. Our results suggested that cell-derived Aβ42 predominantly forms trimers or tetramers, when the Aβ42 size ranged around 13-19 kDa (Figure 3C). These forms have been previously reported to be the most toxic forms of Aβ.^66^ It is not clear whether the observed effect is related only to Aβ42 or to other APP fragments as well present in the conditioned medium. However as mentioned above, only Aβ fragments so far have been confirmed to have the ability to reduce the activity of recombinant HSD10.^10,36–39^ Nevertheless, if other APP fragments still play a role in AD pathology, the reported preparation of cell-derived Aβ42 is an even more physiologically relevant approach when AD samples would contain not only Aβ but also other APP fragments.^67–70^

Motivated by the ability of cell-derived Aβ42 to affect cellular HSD10 activity, the study assessed HSD10 cells’ fitness after cell-derived Aβ42 treatment in a galactose medium. The addition of cell-derived Aβ42 (7.6 nM) decreased ATP levels in combination with elevated signs of cytotoxicity for both HSD10 and HEK293_wt_ treated cell lines in 72 hr (Figure 6B, Supporting Information Figure S16). The 23% decrease in ATP levels and 14% increase in cytotoxicity observed in HEK293_wt_ cells can be considered very similar to the APP_Swe/Ind_ cell line results, albeit these results were not fully comparable because Aβ42 concentration was not quantified inside the APP_Swe/Ind_ cells. Determining Aβ42 concentration inside the cells would be difficult since Aβ42 levels and the rates of Aβ42 secretion can change rapidly, which may complicate accurate measurement of its concentration within the cells.^71^ HSD10 cells exposed to cell-derived Aβ42 displayed an additional 10% reduction in ATP levels accompanied by another 7% increase in cytotoxicity compared with untreated HSD10 cells.

However, the question remains whether this increase in cytotoxicity observed in the HSD10 cells after Aβ42 treatment is simply a sum of the individual cytotoxic effects, or toxicity amplification due to direct interplay between HSD10 and Aβ42, or whether other mechanisms of interplay are involved.

### HSD10 inhibitors do not impair cell fitness in a glucose-free environment

The concept of HSD10 inhibition as a potential treatment of HSD10 overexpression connected with AD pathology was introduced many years ago^10,16,27,72^, and several classes of inhibitors with nanomolar inhibitory ability have been discovered to date.^19,22,24^ To study the cellular HSD10 inhibition in the context of reducing HSD10-driven pathology, two nanomolar HSD10 inhibitors with different mechanisms of inhibition were used: irreversible inhibitor AG18051,^72^ and reversible benzothiazole inhibitor **34**.^73^ In our previous work, both inhibitors were found to be potent for inhibition of recombinant HSD10 using estradiol (E2) assay^23^, having IC_50_ values 89 and 346 nM respectively. Moreover, these inhibitors were shown to penetrate the cells and inhibit the cellular HSD10 enzyme activity along with no observed cytotoxic effects during inhibitor treatment.^23^ Using the fluorogenic probe (-)-CHANA in HSD10 cells we revealed IC_50_ values of 0.19 µM (AG18051) and 4.26 µM (**34**), respectively (Figure 7A). A very similar IC_50_ value for AG18051 inhibitor was previously published confirming the validity of the used methodology.^34^

**Figure 7:**
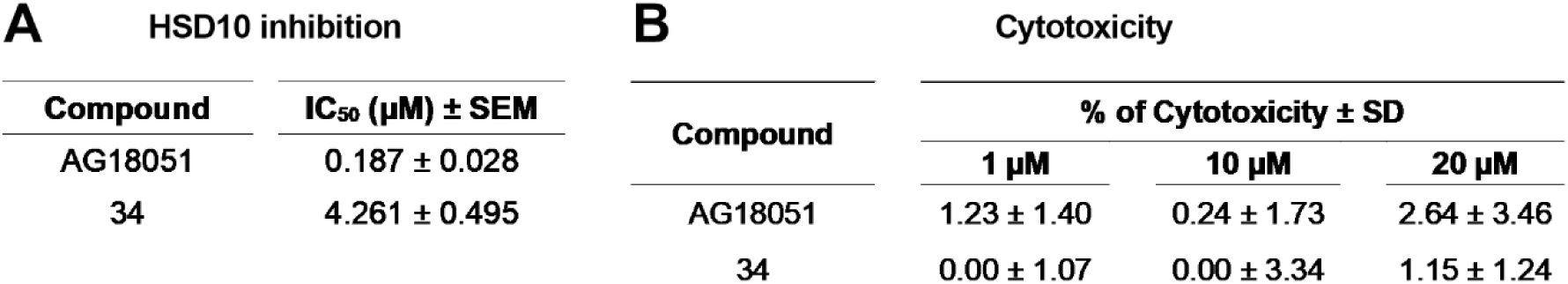
AG18051 and inhibitor **34** characterizations. (**A**) HSD10 inhibition in the cellular environment. The IC_50_ determination of inhibitors AG18051 and **34** using (-)-CHANA fluorogenic probe in HSD10 cells in glucose medium. Values are given as means ± SEM from three cell culture preparations with three technical replicates (n=9). (**B**) Cytotoxicity in HEK293_wt_ after 48 hr treatment of AG18051 or inhibitor **34** (1-20 µM) in galactose medium. Data were normalized between DMSO-treated (1%) and valinomycin-treated (100 µM) HEK293_wt_ cells cultivated in galactose medium. Values are given as means ± SD from two independent cell culture preparations with four technical replicates (n=12). SEM: standard error of the mean. SD: standard deviation.

Since the selected inhibitors have been previously shown to exhibit no cytotoxic effects in the glucose environment^23^, both inhibitors were re-tested in a glucose-free environment to exclude the possibility of hidden cytotoxicity in glucose-free conditions that may bias the results. The cytotoxicity determined 48 hr post-inhibitor treatment in HEK293_wt_ cells in a galactose medium showed no cytotoxicity signs for either inhibitor even at 20 µM concentration (Figure 7B), which indicates their safe use in cell treatments.

### HSD10-connected pathology can be reversed using HSD10 inhibitors

Many of the recently developed HSD10 inhibitors were mostly evaluated on recombinant HSD10 enzymes for their binding and inhibition ability. Several of them were forwarded to cell-based studies, where most of the work was done on AG18051, an irreversible HSD10 inhibitor which forms a covalent adduct with NAD^+^ cofactor within the enzyme’s active site.^72^ This well-characterized inhibitor was used together with our uncompetitive inhibitor **34,** which preferentially binds to enzyme-substrate complex,^73^ to study the potential to reduce the pathological changes associated with HSD10 overexpression. HSD10 cells were cultured in galactose medium, treated with various concentrations of inhibitors (0.5 - 4 equivalents of their IC_50_ values), and assayed after 72 hr incubation.

Both inhibitors (at all four concentrations) caused an increase in the ATP levels and a significant decrease in cytotoxicity of the HSD10 cells (Supporting Information Figure S17). The most potent concentration for both inhibitors was found to be triple the IC_50_ concentration (0.57 µM for AG18051 and 12.78 µM for inhibitor **34**) (Figure 8A, Supporting Information Figure S17), increasing significantly the ATP levels by 18% and reducing the cytotoxicity by almost half. These results confirmed that HSD10 inhibitors can reduce HSD10 overexpression-related pathology. Moreover, the successful reduction of cytotoxic parameters was apparent for both inhibitorś treatments, which differed in the mode of action, and confirmed that the HSD10 inhibition itself, rather than other off-target effects, was responsible for the cell fitness improvement. For AG18051, the related data had already been published in a single study that characterized AG18051 in the setting of HSD10 overexpression alone.^27^ This study was unable to confirm HSD10-induced cytotoxicity but confirmed the disruption of respiratory parameters in HSD10 overexpressing cells, which was restored after AG18051 treatment. Inhibitor **34** has not been previously characterized in the context of HSD10 pathology, and so this is the first report on the cell fitness improvement with reversible and competitive HSD10 inhibitors.

**Figure 8:**
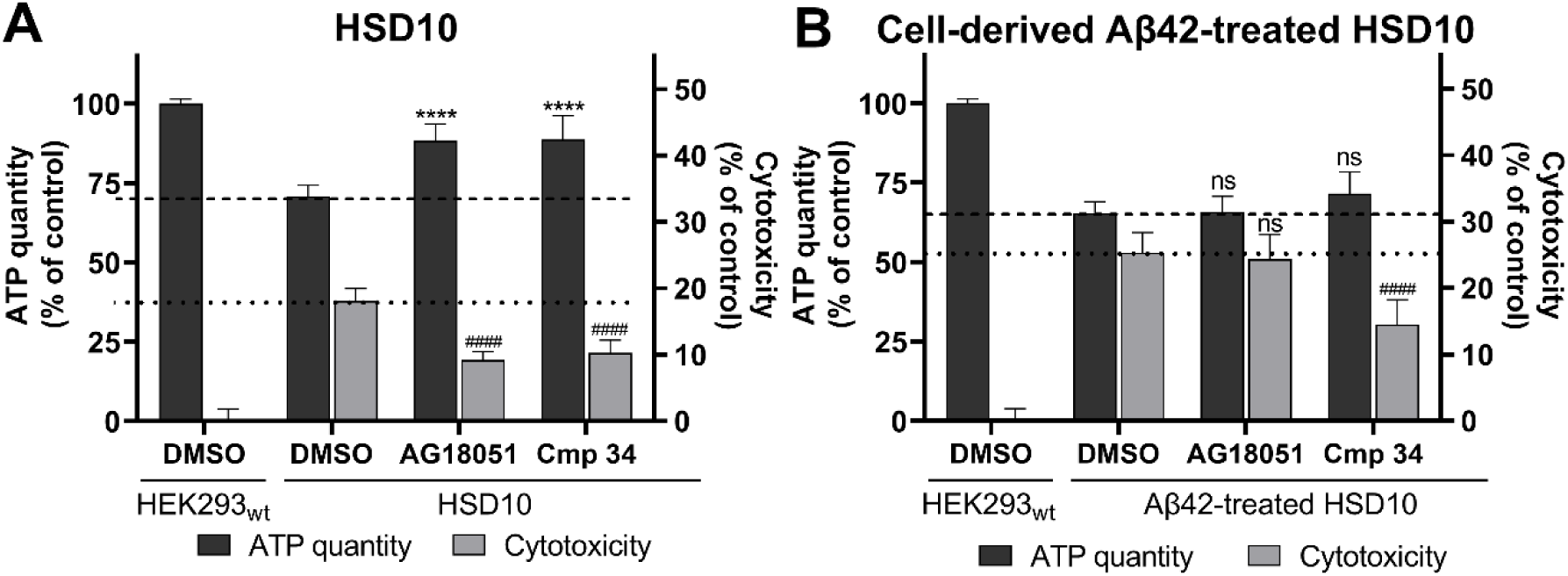
The ability of AG18051 and inhibitor **34** to reduce HSD10-associated cytotoxicity. (**A**) ATP levels and cytotoxicity in HSD10 cells after HSD10-inhibitor treatment performed 72 hr post-seeding into galactose medium. Data were normalized between DMSO-treated (1%) and valinomycin-treated (100 µM) HEK293_wt_ cells cultivated in galactose medium. Values are given as means ± SD from three independent cell culture preparations with four technical replicates (n=12). (**B**) ATP levels and cytotoxicity in HSD10 cells performed 72 hr post-seeding into the Aβ42-containing galactose medium and HSD10-inhibitor treatment. Data were normalized between DMSO-treated (1%) and valinomycin-treated (100 µM) HEK293_wt_ cells cultivated in galactose medium. Values are given as means ± SD from three independent cell culture preparations with three technical replicates (n=9). SD: standard deviation. Statistical difference for ATP quantity: \**p* ≤ 0.05, \*\**p* ≤ 0.01, \*\*\**p* ≤ 0.001, \*\*\*\**p* ≤ 0.0001 compared to the untreated HSD10 control group. Statistical difference for cytotoxicity: ^#^*p* ≤ 0.05, ^##^*p* ≤ 0.01, ^###^*p* ≤ 0.001, ^####^*p* ≤ 0.0001 compared to the untreated HSD10 control group.

### Inhibitor 34 is effective in reducing Aβ42-exacerbated HSD10-driven pathology inside the cells

The Aβ-rich environment and in particular the Aβ42 effect on mitochondria have been repeatedly declared as the determinant of mitochondrial dysfunction and an early pathogenesis trigger leading to neuronal death.^74^ Thus, the therapeutic effect of HSD10 inhibitors was investigated in an Aβ42-rich environment, as it was shown that such environment tends to worsen the observed decline in HSD10 cell fitness (Figure 6B). The ability of HSD10 inhibitors to slow down or even reverse the pathologic phenotype was investigated using HSD10 cells in galactose medium containing 7.6 nM cell-derived Aβ42 (Aβ42 concentration with apparent effect on HSD10 activity) and HSD10 inhibitors AG18051 or **34** in their most potent inhibitory concentrations (3x IC_50_; Figure 8B, Supporting Information Figure S18).

The treatment of HSD10 cells in an Aβ42-rich environment with inhibitor AG18051 had no significant positive effect on improving pathological parameters, with the observed cytotoxicity and ATP production decline remaining completely unchanged upon AG18051 co-administration, a phenomenon confirmed on primary mouse cortical astrocytes.^27^ In contrast, inhibitor **34** resulted in an increase in ATP levels from 65% to 72% and significant cytotoxicity reduction from 25% to 14%, proving its ability to partially reverse the HSD10 pathology in an Aβ42-rich environment. These findings are quite surprising since both inhibitors have been shown to effectively suppress the cytotoxicity of HSD10 overexpression alone (Figure 8A). Based on their confirmed ability to inhibit HSD10 and reverse HSD10-driven pathology, it can be speculated that the ability of **34** to reverse HSD10-induced cytotoxicity in the Aβ42-rich environment might be mediated through molecular targets other than HSD10. As inhibitor **34** is structurally derived from frentizole and Thioflavin-T (a fluorescent Aβ-binding dye) – both known to inhibit the interaction between HSD10 and Aβ^25^ – it raises the question of whether **34** acts through disruption of HSD10-Aβ binding interaction. Alternatively, since inhibitor **34** is an analogue of an Aβ-binding dye with a similar mechanism to that of Thioflavin-T, it may exert its effect through direct binding to Aβ42. These questions need to be further elucidated.

Nevertheless, our data provide a new approach for the evaluation of HSD10 inhibitors in the manner of reducing HSD10-driven pathology either alone or in the presence of Aβ42. The approach presented here allows study of the potential of HSD10 inhibitors to reverse AD-related pathology, suggesting HSD10 inhibitors are beneficial in AD treatment. Concerning AD treatment, the difference in efficacy between AG18051 and **34** exposes the fundamental need for a full understanding of the involved mechanisms, both in the development of the pathology and the mechanisms of action of the inhibitors.

## CONCLUSION

This study was focused on potential pathological triggers which may be causative to AD development, namely HSD10 and APP_Swe/Ind_ overexpressions. Herein, it was confirmed for the first time that HSD10 overexpression itself can be harmful to cell viability in the manner of impaired mitochondrial fitness. The observed cell damage is an outcome of elevated HSD10 enzymatic function: the cellular changes caused by HSD10 overexpression affected several main metabolic pathways, with determination of MEF in the cells indicating decreased activity of TCA cycle enzymes, decreased β-oxidation of fatty acids, and increased oxidative stress. It is suggested that these metabolic changes are likely related to a decrease in E2 levels as a consequence of increased HSD10 activity. These findings provide new insight into HSD10 overexpression in AD, and suggest that HSD10 may not simply serve as an exacerbating factor to Aβ-driven damage but have its own unique role in the pathology. Additionally, the APP_Swe/Ind_ overexpression and associated Aβ42 production were shown to also induce cell toxicity via mitochondrial damage. Unlike the HSD10 cell line, the observed metabolic changes exposed by the MEF determination via the Mitoplate S-1 assay suggest that APP_Swe/Ind_ overexpression resulted in glucose hypermetabolism, associated upregulation of β-oxidation, and changes in the TCA cycle. These data correlate with previous findings suggesting an increase in glycolysis as a protective mechanism against mitochondrial damage caused by Aβ production in the early stages of AD.^44,62^ In addition, the pathologies induced by the studied overexpressions were mainly manifested in a glucose-free environment, confirming the previous suggestions that problems in glucose utilization may serve as an important factor responsible for the development of neuronal defects observed in AD.^75^

Moreover, the use of APP_Swe/Ind_ conditioned media as a source of Aβ42 was established for use in other *in vitro* experiments. The cell-derived Aβ42 displayed the ability to affect HSD10 enzymatic activity and further potentiate the HSD10-driven cellular damage. Although the precise mechanism by which HSD10 and Aβ42 crosstalk affects cellular fitness remains unclear, the metabolically distinct basis of HSD10 and APP_Swe/Ind_ overexpression indicates that other mechanisms beyond direct HSD10-Aβ42 binding may also be involved. Last but not least, a new opportunity was presented to evaluate HSD10 inhibitors in the context of their potential to reduce HSD10-driven pathology, both in the presence and absence of Aβ42, thus advancing the research and development of HSD10 inhibitors. This newly developed cell-based assay was used to characterize two known HSD10 inhibitors. Data revealed that HSD10 inhibitors have the ability to reverse HSD10-related pathology and highlighted the benzothiazole inhibitor **34** as a promising AD drug candidate efficient in restoring cell viability and decreasing cytotoxicity even in the Aβ42-rich environment, and thus be effective in amyloidogenic stress suppression.

## MATERIALS AND METHODS

### Chemicals and compounds

The majority of the chemicals used were obtained from Sigma-Aldrich unless otherwise noted. All chemicals were of the highest commercially available purity. The tested HSD10 inhibitors were chosen from the in-house library of the University of Hradec Kralove, Faculty of Science. The synthetic routes and chemical properties can be found in the published articles.^72,73^ All compounds were dissolved in anhydrous dimethyl sulfoxide (DMSO) and further diluted with assay buffers or culture media to the working concentrations to keep maximum DMSO concentration at 2% (v/v) for HSD10 activity measurements and at 1% (v/v) for other cell culture experiments.

### Cell cultures and HSD10 overexpression or APP_Swe/Ind_ overexpression

HEK293 cells (ECACC Cat# 85120602, RRID: CVCL_0045) were commonly cultured in Dulbeccós modified Eagle medium (Capricorn; standard DMEM medium with phenol red) supplemented with 10% foetal bovine serum (Gibco), 2 mM L-glutamine (Gibco) and non-essential amino acid additives (Gibco) at 37 °C in 6% CO_2_ humidified atmosphere (standard DMEM medium). The cells were passaged regularly at 70-80% confluency and routinely tested for *Mycoplasma* contamination (MycoAlert Plus, Promega). All experiments were performed at a maximum passage number of 15 (the passage number counting started at the time of purchase). Glucose (4.5 g/L) or galactose (1.8 g/L) Dulbecco’s modified Eagle medium (Gibco, A1443001) without the addition of phenol red supplemented with 0.11 g/L sodium pyruvate, 10% foetal bovine serum (Gibco), 2 mM L-glutamine and non-essential amino acid additives (Gibco) were used for all cell culture experiments. Expi293 Expression Medium (Gibco, A1435101) was used for Western blot analysis, size-exclusion chromatography, and mass-spectrometry analysis of the conditioned culture medium.

For APP_Swe/Ind_ overexpression, pCAX APP695 Swe/Ind vector (pCAX APP Swe/Ind was a gift from Dennis Selkoe^76^ & Tracy Young-Pearse (Addgene plasmid # 30145; http://n2t.net/addgene:30145; RRID: Addgene_30145)) was used as a template. The full-length gene for APP_Swe/Ind_ was PCR amplified and inserted into constitutive mammalian expression pcDNA3.4 vector using topoisomerase-based cloning and the insertion was subsequently sequenced. Low-passaged wild-type HEK293 cells (HEK293_wt_) were nucleofected (Amaxa Nucleofector kit V, Lonza, nucleofection program Q-001) with 3 µg of pcDNA3.4-APP_Swe/Ind_ vector plasmid DNA isolated using PureLink HiPure Plasmid MiniPrep kit (Invitrogen). Twenty-four hours after nucleofection, the cells were treated with G418 disulphate salt solution (Roche) at a concentration of 500 µg/mL in culture medium to select positive cell clones. After several weeks of clonal selection and expansion, several monoclonal cell lines (referred to as APP_Swe/Ind_ cells) were isolated using the limiting dilution method. The APP_Swe/Ind_ overexpression and subsequent Aβ42 overproduction were confirmed by immunoblotting using anti-APP and anti-Aβ primary rabbit monoclonal antibodies (Table 1). For HSD10 overexpression, HSD10 overexpressing cells published previously were used.^23^ For the preparation of HSD10 overexpression cell lines harbouring the inactivated catalytic active site of the enzyme (Y168A, K172A) (referred to as HSD10_mut_ cells), the HSD10 DNA was modified accordingly, prepared *de novo* as a DNA string (GeneArt, ThermoFisher) and cloned into pcDNA3.4 vector. After DNA sequencing verification, the same procedure was followed as for APP_Swe/Ind_ cell line preparation. The final confirmation of the expression level was performed using an anti-ERAB monoclonal antibody (Table 1). No institutional ethical approval was required for this work.

**Table 1:**
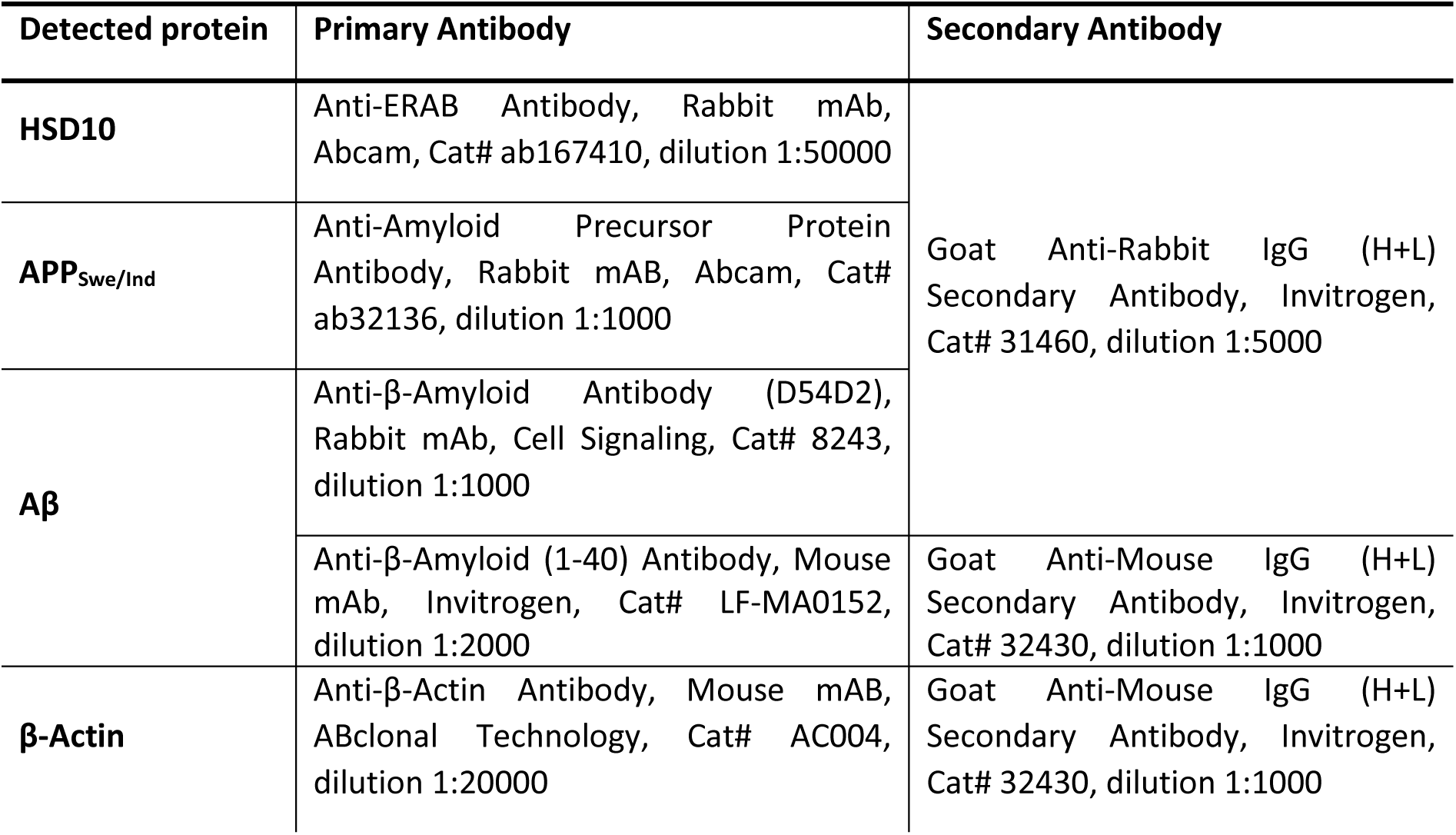
Primary and secondary antibodies used for western blot analysis.

| Detected protein | Primary Antibody | Secondary Antibody |
| --- | --- | --- |
| **HSD10** | Anti-ERAB Antibody, Rabbit mAb, Abcam, Cat# ab167410, dilution 1:50000 | Goat Anti-Rabbit IgG (H+L) Secondary Antibody, Invitrogen, Cat# 31460, dilution 1:5000 |
| **APP<sub>Swe/Ind</sub>** | Anti-Amyloid Precursor Protein Antibody, Rabbit mAb, Abcam, Cat# ab32136, dilution 1:1000 |  |
| **Aβ** | Anti-β-Amyloid Antibody (D54D2), Rabbit mAb, Cell Signaling, Cat# 8243, dilution 1:1000 | |
| | Anti-β-Amyloid (1-40) Antibody, Mouse mAb, Invitrogen, Cat# LF-MA0152, dilution 1:2000 | Goat Anti-Mouse IgG (H+L) Secondary Antibody, Invitrogen, Cat# 32430, dilution 1:1000 |
| **β-Actin** | Anti-β-Actin Antibody, Mouse mAb, ABclonal Technology, Cat# AC004, dilution 1:20000 | Goat Anti-Mouse IgG (H+L) Secondary Antibody, Invitrogen, Cat# 32430, dilution 1:1000 |

### Western blot analysis

The expression levels of particular proteins in the overexpressed cell lines were confirmed using sodium dodecyl sulfate-polyacrylamide gel electrophoresis 10% (APP_Swe/Ind_ detection) or 15% (HSD10, HSD10_mut_ and Aβ detection) resolving gel and 4% stacking gel. The protein samples under reducing conditions were subjected to electrophoresis (10 µg/20 µg of whole cell lysates from HEK293_wt_, HSD10, HSD10_mut,_ or APP_Swe/Ind_ cell lines or 14 µL of conditioned culture media from HEK293_wt_ or APP_Swe/Ind_ lines), subsequently transferred to 0.45 µm PVDF membrane, and detected using a primary antibody followed by chemiluminescence detection of HRP-conjugated secondary antibody (Amersham ECL Prime). β-actin as a loading control was used for analysis of whole cell lysates. The gel staining was used as a loading control for the conditioned medium analysis. Primary and secondary antibodies are listed in Table 1.

### ATP level and cytotoxicity determination

The impact of overexpression of particular proteins (HSD10, HSD10_mut_ or APP_Swe/Ind_) or of HSD10 inhibitors on ATP production and the potential cytotoxic effects was tested using CellTiter-Glo Luminescent Cell Viability Assay and CellTox Green Cytotoxicity Assay kits (Promega, G7574 and G8741), respectively. For multiplex measurement, 5×10^3^ cells were seeded in 50 µL of glucose or galactose culture medium per well into a white 96-well microplate with a clear flat bottom (Corning, CLS3903). Cells were cultured for 24, 72, or 168 hr depending on the experiment setup. The assay readouts were measured using the TECAN Spark 10 M instrument following the manufacturer’s protocol with minor changes. For cytotoxicity determination, 20 µL of complete CellTox Green Reagent (mixing 20 µL of CellTox Green Dye with 6 mL of CellTox Green Assay Buffer) was used in combination with fluorescence readout (Ex/Em=485/530 nm). For ATP production measurements, 25 µL of CellTiter-Glo 2.0 Reagent was used, followed by the luminescence-based measurement with an integration time of 500 ms. HEK293_wt_ cells treated with 1% DMSO only (vehicle control), and 100 µM valinomycin-treated cells (positive control) were used as the controls.

### Viability determination

The protein overexpression effect on cell viability was measured using RealTime-Glo MT Cell Viability Assay (Promega, G9711). The cells (5×10^3^) were seeded in 50 µL of complete glucose medium per well into a white 96-well microplate with a clear flat bottom (Corning, CLS3903). Cells were cultured for 24 hr to allow them to adhere, followed by changing the culture medium to galactose, immediately treating with 50 µL of galactose medium supplemented with MT Cell Viability Substrate and NanoLuc Enzyme in final 2X concentration. The treated cells were incubated at 37°C and 6% CO_2_ for 30 min followed by 72 hr of luminescence-based viability monitoring (every 6 or 12 hr) using a Tecan Spark 10 M instrument.

### Mitochondrial toxicity determination

A Mitochondrial ToxGlo Assay kit (Promega, G8000) was used to determine the effect of protein overexpression (HSD10 or APP_Swe/Ind_) on mitochondrial fitness. For measurement, 5×10^3^ cells were seeded in 50 µL complete galactose medium per well into a white 96-well microplate with a clear flat bottom (Corning, CLS3903) followed by 72 hr incubation time. Several control treatments were performed by adding 50 μL of control-containing media for different time intervals: positive toxicity treatment (100 µM valinomycin treated HEK293_wt_ cells) for 24 hr; positive mitochondrial toxicity treatment (1 µM antimycin A treated HEK293_wt_ cells) for 90 min; and vehicle control (1% DMSO treated HEK293_wt_ or overexpressing cells) for 90 min. The mitochondrial dysfunction was determined as the activity of dead-cell proteases in combination with ATP level measurement. Complete Cytotoxicity Reagent treatment (20 µL; mixing of 10 µL bis-AAF-R110 substrate with 2 mL Mitochondrial ToxGlo Assay Buffer) was added to the cells for 30 min and measured fluorometrically (Ex/Em=485/530 nm, Tecan Spark 10 M), followed immediately by treatment with 100 μL 2x ATP Detection reagent and luminescence measurement of ATP levels with an integration time of 500 msec.

### Aβ42 quantification

To quantify the amount of Aβ42 secreted by the APP_Swe/Ind_ cell line, the Amyloid beta 1-42 Kit (CisBio, 62B42PEG) was used. After 3 to 5 days of cultivation of APP_Swe/Ind_ cells, the glucose or galactose-conditioned medium was collected and centrifuged at 10,000 RCF for 10 min to clear the supernatant. The reaction mixture consisted of an HTRF binding pair of donor and acceptor antibodies (the Human Amyloid β 1-42 Europium cryptate antibody, and the Human Amyloid β 1-42 d2 antibody diluted with Detection buffer to 1X concentration, 2 μL each) and 16 μL of diluted sample medium (20 to 50X) or Aβ42 standard. The assay mixture was pipetted into an HTRF 96-well low-volume white plate (Revvity, 66PL96005), sealed, and incubated overnight at RT. Two sequential measurements of the FRET signal at 620 nm for Eu-cryptate emission (donor) and 665 nm for the d2 emission (acceptor) were measured followed by calculation of the 665/620 ratio that was used for concentration determination based on the standard curve approximation.

### Cellular HSD10 inhibition

HSD10 activity in the cellular environment was performed using (-)-CHANA (cyclohexenyl amino naphthalene alcohol) fluorogenic probe.^23,24,34^ The cells were seeded at a density of 1×10^4^ cells per well in 200 µL of complete glucose medium into a black 96-well microplate with a clear bottom (Brand, BR781971). After 20 hr of incubation, the cells were treated with DMSO only, Aβ42 containing medium, or HSD10 inhibitors dissolved in anhydrous DMSO. After 2 hr of treatment, 2 µL of (-)-CHANA probe was added at the final concentration of 20 µM. Changes in fluorescent intensities were measured immediately (0 hr) and after 2 hr of incubation. The fluorescence intensities of the reaction product cyclohexenyl amino naphthalene ketone (CHANK) were measured using the TECAN Spark 10 M instrument (Ex/Em=380/525nm). The HSD10 activity was calculated as the ΔF between 2 hr and 0 hr after (-)-CHANA addition and the data were normalized between DMSO-treated HSD10 cells and non-transfected HEK293_wt_ controls (using relative response ratio).

### Cell metabolic activity changes

To investigate the changes in the cell metabolic activities caused by HSD10 or APP_Swe/Ind_ overexpression, the MitoPlate S-1 technology (Biolog) was used. The 96-well plates preloaded with 31 different substrates (TCA cycle intermediates, amino acids, fatty acids, hexoses, and trioses, each in triplicate) were activated by adding 30 µL of Assay Mix (15 µl 2x Biolog Mitochondrial Assay Solution; 10 µL 6x Redox Dye MC; 2.5 µL 24x saponin (Sigma, #47036) at 960 µg/mL, final concentration 40 µg/mL; and 2.5 µL sterile water) per well and incubated for 1 hr at 37 °C and 6% CO_2_. The cells were cultivated for 48 hr before the experiment in glucose or galactose complete media and then plated into the activated MitoPlate S-1 plates at a density of 3×10^4^ per well in 30 µL of 1X Biolog Mitochondrial Assay Solution. The reduction of the Redox Dye mix, which corresponds to changes in mitochondrial electron flow, was read by absorbance (OD_590_) kinetically on Tecan Spark 10 M instrument at 30-sec intervals for a total of 4 hr after seeding. The data was extracted as ΔAbs per hour from the linear phase of the particular substrate consumption rates.

### Liquid chromatography-mass spectrometry analysis

APP_Swe/Ind_ conditioned cell culture medium was loaded to DSC-18 500 mg/mL SPE (Sigma) and bound material was eluted with 20% acetonitrile (ACN) and lyophilized. The lyophilizate was dissolved in reducing LDS buffer (Invitrogen), heated at 70 °C, divided into four lanes of NuPAGE precast gel (Invitrogen), and run by SDS-PAGE. The gel was stained and visible bands were excised, chopped, and combined from each lane. Protein digestion was done as previously described.^77^ Peptides were analyzed using the UltiMate 3000 RSLCnano system connected with Orbitrap Exploris 480 equipped with FAIMS interface (Thermo Fisher Scientific). Samples were separated on analytical column (PepMap RSLC C18; 75 µm×250 mm; Thermo Fisher Scientific) by running a linear gradient of phase B (80% ACN/0.1% FA) over phase A (2% ACN/0.1% FA). MS analysis was done using data-dependent acquisition with selected precursors fragmented by HCD. Sample material from <15 kDa band was further re-analyzed by a longer LC-MS method. Data were interpreted using MSFragger (v4.1)^78^. Methionine oxidation and N-terminal acetylation were set as variable and carbamidomethylation of cysteine was set as a fixed modification. Data were searched against the Human Reference proteome database (UniProt) combined with sequences of APP_Swe/Ind_ variant and Aβ fragment. Proteins were filtered at 1% FDR.

### Size-exclusion chromatography

The molecular weight of the Aβ42 peptide derived from APP_Swe/Ind_ conditioned medium was determined by size-exclusion chromatography using an NGC Medium-Pressure Chromatography System (Bio-Rad, USA) equipped with a Superdex 200, 10/300 GL column (Cytiva, USA). Phosphate-buffered saline (PBS, pH 7.4) was used as the mobile phase, using the 0.7 mL/min flow rate and 0.35 mL injection volume. The samples were separated into 28 fractions followed by immunoblotting detection using an anti-Aβ antibody. The fractions with positive antibody detection were evaluated based on a calibration curve generated using a Cytiva low molecular weight gel filtration calibration kit (28403841).

### Statistical analysis

Data were calculated as mean ± SD from single experiments. Statistical analyses were performed using GraphPad Prism 9 software (GraphPad Software, San Diego, CA). Ordinary one-way ANOVA with Tukey’s post hoc test was employed and *p-*value < 0.05 was considered statistically significant.

## SUPPORTING INFORMATION

Original western blot images, full data from cell-based methods, and liquid chromatography-mass spectrometry analysis are available in **Supporting Information**.

## AUTHOR CONTRIBUTIONS

A.H. and M.S. designed the study; A.H., M.S., I.F., and R.A. performed experiments; A.H. and M.S. wrote the manuscript, A.H., M.S., O.B., I.F., L.Z, O.S., and K.M. revised the manuscript

## ACKNOWLEDGMENT

This work was supported by the University of Hradec Kralove (Faculty of Science, no. *SV2112/2024*), MH CZ – DRO (UHHK, 00179906), and The Biomedical Indicators for Personalized Medicine project (BIPOLE, ID CZ.02.01.01/00/23_021/0008439), which is co-funded by the European Union. The authors are grateful to Ian McColl MD, PhD for assistance with the manuscript.

## ABBREVIATION

Aβ: amyloid-beta peptide
ACN: acetonitrile
AD: Alzheimer’s disease
APP: amyloid precursor protein
ATP: adenosine triphosphate
C99: C-terminal 99 amino acid long fragment of amyloid precursor protein
CHANA: cyclohexenyl amino naphthalene alcohol
CHANK: cyclohexenyl amino naphthalene ketone
DMEM: Dulbecco’s Modified Eagle Medium
DMSO: dimethyl sulfoxide
DTT: dithiothreitol
E2: 17β-estradiol
ECL: excellent chemiluminescent substrate
ERAB: endoplasmic reticulum-associated amyloid β-peptide-binding protein
FA: formic acid
FADH_2_/FAD: flavin adenine dinucleotide
HEK293: human embryonic kidney cells
HK: hexokinase
HSD10: 17β-hydroxysteroid dehydrogenase type 10
LC-MS: liquid chromatography-mass spectrometry
MAO: monoamine oxidase
MOPS: 3-(*N*-morpholino)propanesulphonic acid
MS/MS: tandem mass spectrometry
NADH/NAD^+^: nicotinamide adenine dinucleotide
OXPHOS: oxidative phosphorylation
PCR: polymerase chain reaction
RCF: relative centrifugal force
ROS: reactive oxygen species
RNase P: ribonuclease P
RNS: reactive nitrogen species
sAPPβ: N-terminal cleavage fragment of amyloid precursor protein
SD: standard deviation
SDS-PAGE: sodium dodecyl-sulphate polyacrylamide gel electrophoresis
SEM: standard error of the mean
TCA: tricarboxylic acid cycle
TEAB: triethylammonium bicarbonate buffer
wt: wild-type

