## Supporting Information for "New insights into the 17β-hydroxysteroid dehydrogenase type 10 and amyloid-β 42 derived cytotoxicity relevant to Alzheimer’s disease"

**TABLE OF CONTENTS:**

1. Appendix Figure S1: Immunoblotting analysis of HSD10, and $HSD10_{mut}$ cell (related to Figure 1A).
2. Appendix Figure S2: Immunoblotting analysis of HSD10, and $HSD10_{mut}$ cell (related to Figure 1A).
3. Appendix Table S3: HSD10 activity determination in $HEK293_{wt}$, HSD10, $HSD10_{mut}$, and $APP_{Swe/Ind}$ cells (related to Figures 1B and 2D).
4. Appendix Figure S4: Immunoblotting analysis of $APP_{Swe/Ind}$ cells (related to Figure 2A).
5. Appendix Figure S5: Immunoblotting analysis of $APP_{Swe/Ind}$ cells (related to Figure 2A).
6. Appendix Figure S6: Immunoblotting analysis of $APP_{Swe/Ind}$ cells (related to Figure 2B).
7. Appendix Figure S7: Immunoblotting analysis of $APP_{Swe/Ind}$ conditioned medium (related to Figure 2C).
8. Appendix Figure S8: Immunoblotting analysis of $APP_{Swe/Ind}$ conditioned medium (related to Figure 2C).
9. Appendix Figure S9: Analysis of $APP_{Swe/Ind}$ conditioned medium (related to Figures 3A and 3B).
10. Appendix Figure S10: Immunoblotting analysis of $APP_{Swe/Ind}$ conditioned medium fractions (related to Figure 3C).
11. Appendix Table S11: Chromatogram of standards' separation by size-exclusion chromatography (related to Figure 3C).
12. Appendix Table S12: ATP levels and cytotoxicity in $HEK293_{wt}$, HSD10, $HSD10_{mut}$, and $APP_{Swe/Ind}$ cells (related to Figures 4A and 4B).
13. Appendix Table S13: Viability of $HEK293_{wt}$, HSD10, $HSD10_{mut}$, and $APP_{Swe/Ind}$ cells (related to Figures 4C).
14. Appendix Table S14: Mitochondrial toxicity of HEK293, HSD10, and $APP_{Swe/Ind}$ cells (related to Figures 4D).
15. Appendix Table S15: Mitochondrial electron flow changes (correspond to metabolic changes) in $HEK293_{wt}$, HSD10, and $APP_{Swe/Ind}$ cells (related to Figure 5).
16. Appendix Table S16: ATP levels and cytotoxicity in $HEK293_{wt}$, and HSD10 cells after cell-produced Aβ42 treatment (related to Figure 6B).

17. Appendix Table S17: ATP levels and cytotoxicity in HSD10 cells after HSD10-inhibitors treatment (related to Figure 8A).
18. Appendix Table S18: ATP levels and cytotoxicity in HSD10 cells after HSD10-inhibitors and cell-produced Aβ42 treatment (related to Figure 8B).

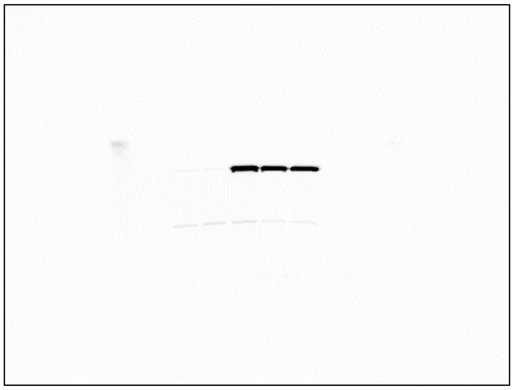

**Figure S1**: Original western blot image from $HEK293_{wt}$, HSD10, and $HSD10_{mut}$ cell lysates immunoblotting analysis using an anti-HSD10 primary antibody.

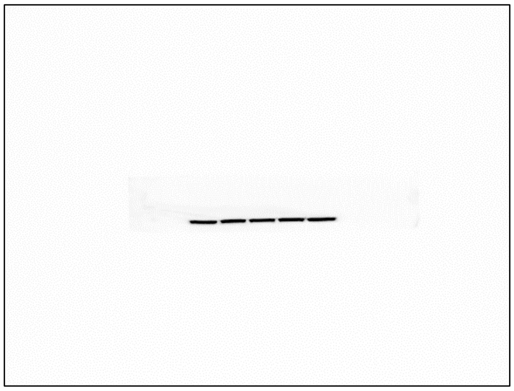

**Figure S2:** Original western blot image from $HEK293_{wt}$, HSD10, and $HSD10_{mut}$ cell lysates immunoblotting analysis using an anti-β-Actin primary antibody.

**Table S3:** HSD10 activity determination via (-)-CHANA to CHANK turnover in $HEK293_{wt}$, HSD10, $HSD10_{mut}$, and $APP_{Swe/Ind}$ cells.

| Cell line | ΔF (2 hr) |
|---|---|
| **$HEK293_{wt}$** | 77.26 ± 83.61 |
| **HSD10** | 1050.41 ± 73.71 |
| **$HSD10_{mut}$** | 84.37 ± 79.48 |
| **$APP_{Swe/Ind}$** | -210.14 ± 95.52 |

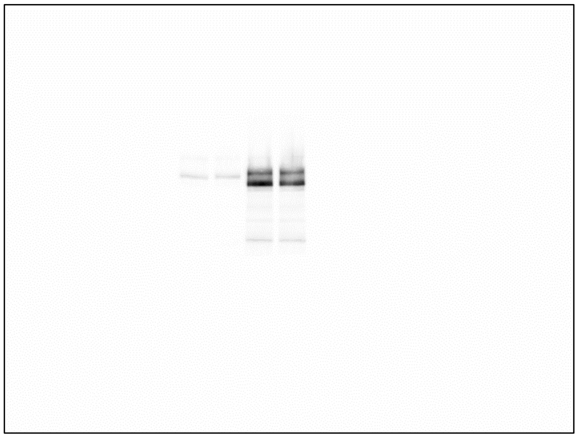

**Figure S4**: Original western blot image from HEK293$_{wt}$ and APP$_{Swe/Ind}$ cell lysates immunoblotting analysis using an anti-APP primary antibody.

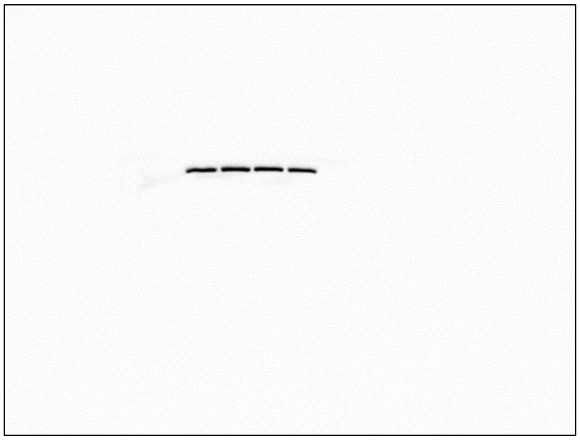

**Figure S5:** Original western blot image from HEK293$_{wt}$ and APP$_{Swe/Ind}$ cell lysates immunoblotting analysis using an anti-β-Actin primary antibody.

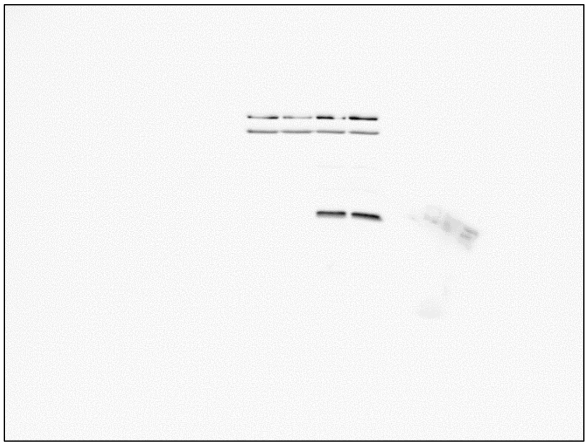

**Figure S6:** Original western blot image from $HEK293_{wt}$ and $APP_{Swe/Ind}$ cell lysates immunoblotting analysis using a combination of anti-Aβ, and anti-β-actin primary antibodies.

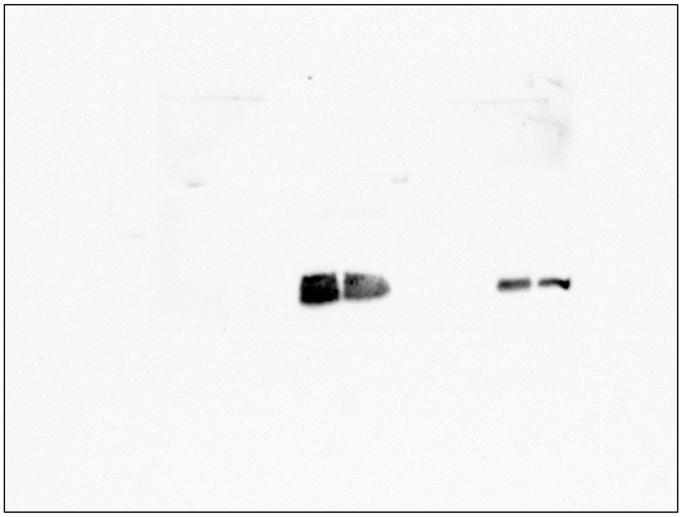

**Figure S7:** Original western blot image from $HEK293_{wt}$ and $APP_{Swe/Ind}$ cell conditioned medium immunoblotting analysis using an anti-Aβ primary antibody.

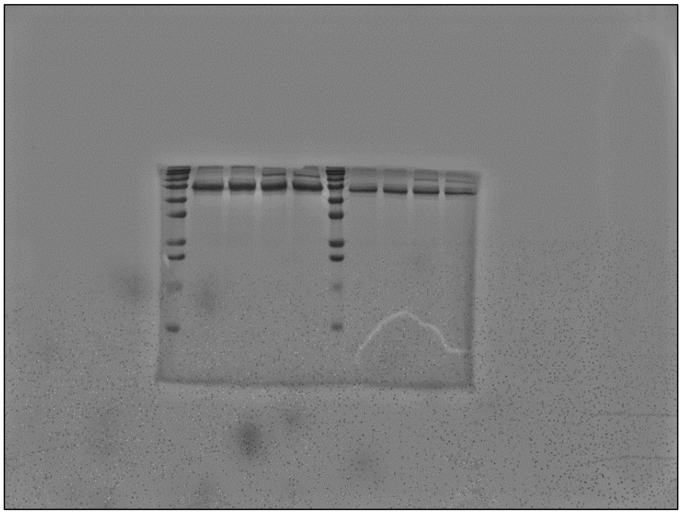

**Figure S8:** Original gel staining image from HEK293$_{wt}$ and APP$_{Swe/Ind}$ cells conditioned medium immunoblotting analysis.

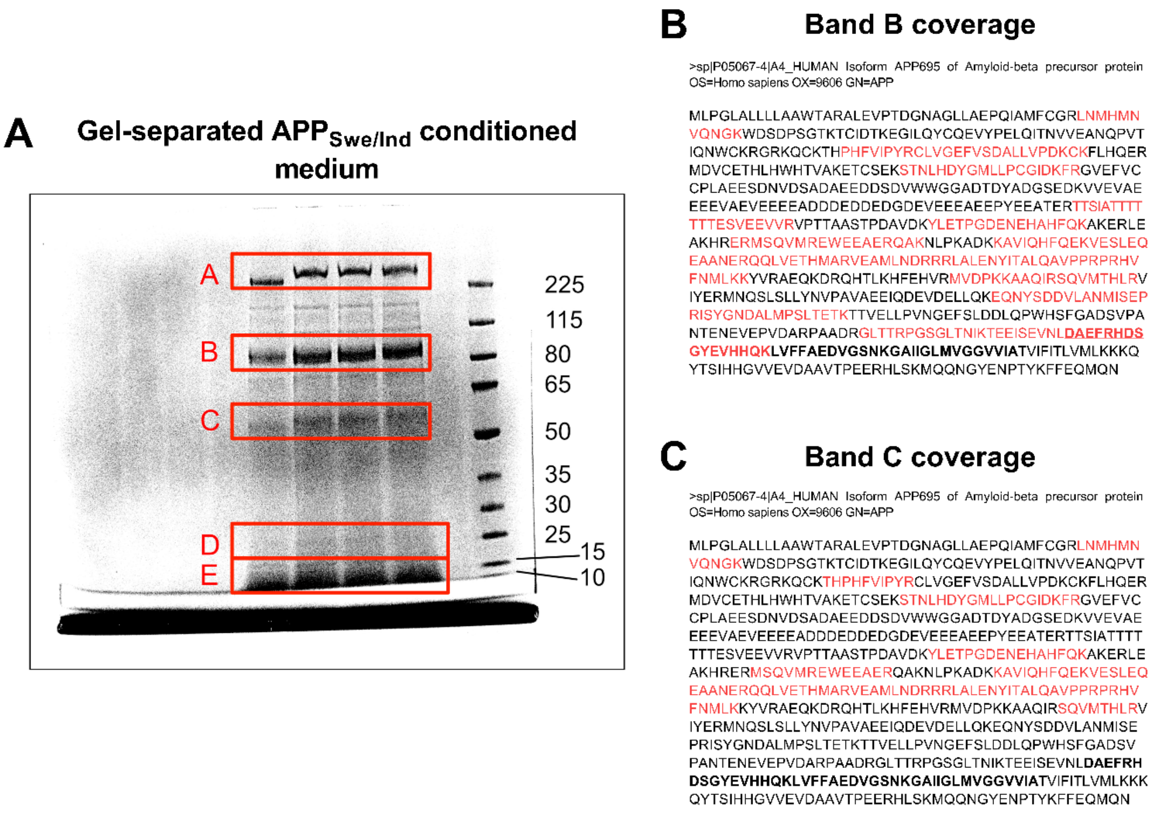


**Figure S9:** Analysis of APP$_{Swe/Ind}$ conditioned medium. (**A**) The APP$_{Swe/Ind}$ conditioned serum and protein-free medium was desalted, and column separated, individual fractions were lyophilized, and gel-separated in four lanes of NuPAGE gel. Visible bands from the concentrated samples were excised and subjected to mass-spectrometric analysis, which revealed the presence of APP or APP-derived fragments in Band B, C, and E. (**B**)

Coverage (marked in red) of tryptically digested proteins from Band B with the FASTA sequence of $APP_{Swe/Ind}$. The results suggest the presence of full-length $APP_{Swe/Ind}$ protein. (**C**) Coverage (marked in red) of tryptically digested proteins from Band C with the FASTA sequence of $APP_{Swe/Ind}$. The results suggest the presence of APP-derived N-terminal cleavage fragments.

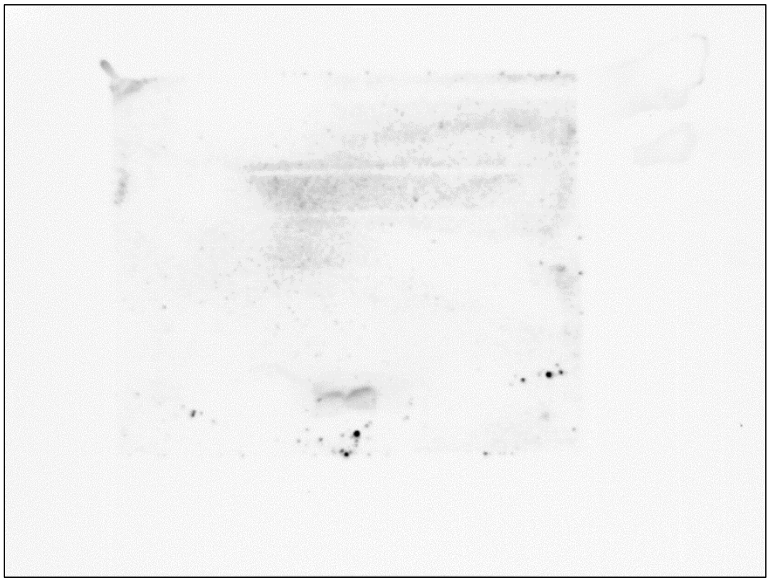

**Figure S10:** Original western blot image from $APP_{Swe/Ind}$ conditioned medium fractions immunoblotting analysis using an anti-Aβ primary antibody.

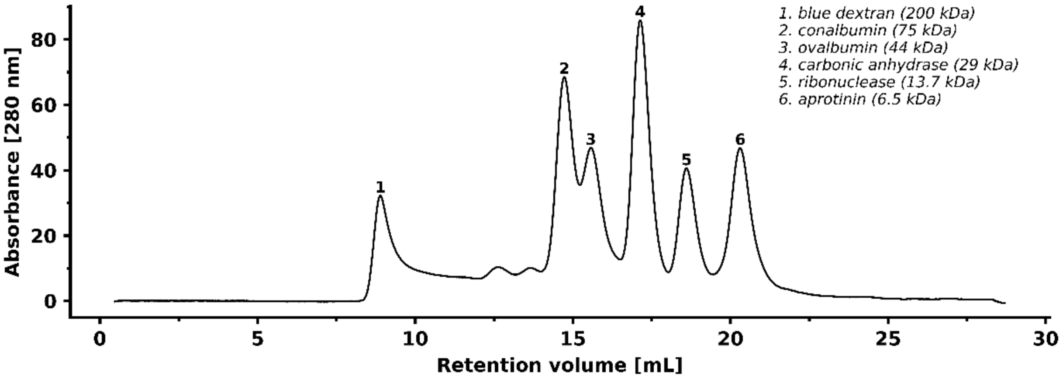


**Figure S11:** Size-exclusion chromatogram of the Cytiva low molecular weight gel filtration calibration kit (28403841) using a Superdex 75 10/300 GL column. Mobile phase: PBS, flow rate: 0.7 mL/min, detection at 280 nm. Peaks are labeled with their corresponding molecular weights (kDa).

**Table S12:** ATP levels and cytotoxicity in $HEK293_{wt}$, HSD10, $HSD10_{mut}$, and $APP_{Swe/Ind}$ cells 72 hr and 168 hr post-seeding into the glucose and galactose medium. Data were normalized between DMSO-treated (1%) and valinomycin-treated (100 µM) $HEK293_{wt}$ cells cultivated in glucose media. Values are given as means ± SD from three independent cell culture preparations with four technical replicates (n=12). SD; standard deviation.

| **Cell line** | **72 hr** | | | | **168 hr** | | | |
|---|---|---|---|---|---|---|---|---|
| | **Glucose** | | **Galactose** | | **Glucose** | | **Galactose** | |
| | **% of ATP quantity** | **% of Cytotoxicity** | **% of ATP quantity** | **% of Cytotoxicity** | **% of ATP quantity** | **% of Cytotoxicity** | **% of ATP quantity** | **% of Cytotoxicity** |
| **$HEK293_{wt}$** | 100.00 ± 2.33 | 0.00 ± 1.65 | 89.45 ± 3.84 | 2.86 ± 2.61 | 100.00 ± 2.52 | 0.00 ± 1.45 | 87.05 ± 3.93 | 4.17 ± 2.70 |
| **HSD10** | 76.67 ± 3.09 | 1.81 ± 1.67 | 69.23 ± 2.78 | 14.63 ± 3.09 | 71.75 ± 6.53 | 4.52 ± 3.64 | 50.11 ± 4.42 | 39.33 ± 5.09 |
| **$HSD10_{mut}$** | 93.98 ± 8.66 | 1.64 ± 0.78 | 84.49 ± 4.82 | 3.76 ± 2.33 | 92.07 ± 4.29 | 3.36 ± 1.07 | 82.46 ± 4.12 | 5.58 ± 1.45 |
| **$APP_{Swe/Ind}$** | 66.37 ± 3.98 | 3.06 ± 2.53 | 54.05 ± 4.35 | 11.33 ± 2.25 | 63.18 ± 5.19 | 5.33 ± 3.60 | 46.82 ± 4.35 | 37.73 ± 3.37 |

**Table S13:** Viability of $HEK293_{wt}$, HSD10, $HSD10_{mut}$, and $APP_{Swe/Ind}$ cells monitored for 72 hr of galactose media cultivation. Values are given as means ± SD from three independent cell culture preparations with three technical replicates (n=9). SD; standard deviation.

| Cell line | Viability (RLU) | | | | | | | | | |
|---|---|---|---|---|---|---|---|---|---|---|
| | **0 hr** | **6 hr** | **12 hr** | **24 hr** | **30 hr** | **36 hr** | **48 hr** | **54 hr** | **60 hr** | **72 hr** |
| **$HEK293_{wt}$** | 12174 ± 1179 | 27338 ± 1684 | 40672 ± 3151 | 59211 ± 4932 | 67956 ± 2926 | 75202 ± 2529 | 86520 ± 1828 | 93972 ± 2230 | 104795 ± 3664 | 121752 ± 6226 |
| **HSD10** | 9733 ± 303 | 22567 ± 899 | 33489 ± 1151 | 52152 ± 1950 | 57042 ± 2181 | 63138 ± 2641 | 67950 ± 2408 | 71303 ± 3146 | 75670 ± 1877 | 82197 ± 4629 |
| **$HSD10_{mut}$** | 12057 ± 988 | 23385 ± 2085 | 36381 ± 1996 | 52922 ± 4330 | 60904 ± 4668 | 70349 ± 2816 | 79650 ± 3233 | 87653 ± 2744 | 98823 ± 3299 | 109114 ± 2935 |
| **$APP_{Swe/Ind}$** | 7877 ± 960 | 20243 ± 721 | 28929 ± 1043 | 44196 ± 1847 | 48473 ± 2210 | 55747 ± 2857 | 64082 ± 3632 | 70102 ± 3555 | 76341 ± 3276 | 82372 ± 2828 |

**Table S14:** Mitochondrial toxicity of HEK293, HSD10, and $APP_{Swe/Ind}$ cells 72 hr post-seeding to galactose media. Values are given as means ± SD from three independent cell culture preparations with four technical replicates (n=12). SD; standard deviation.

| Cell line | % of ATP levels | % of Dead-cell protease activity |
|---|---|---|
| **$HEK293_{wt}$** | 100.00 ± 4.61 | 0.00 ± 0.13 |
| **HSD10** | 59.20 ± 2.32 | 2.61 ± 0.83 |
| **$APP_{Swe/Ind}$** | 56.82 ± 2.74 | 3.42 ± 0.78 |

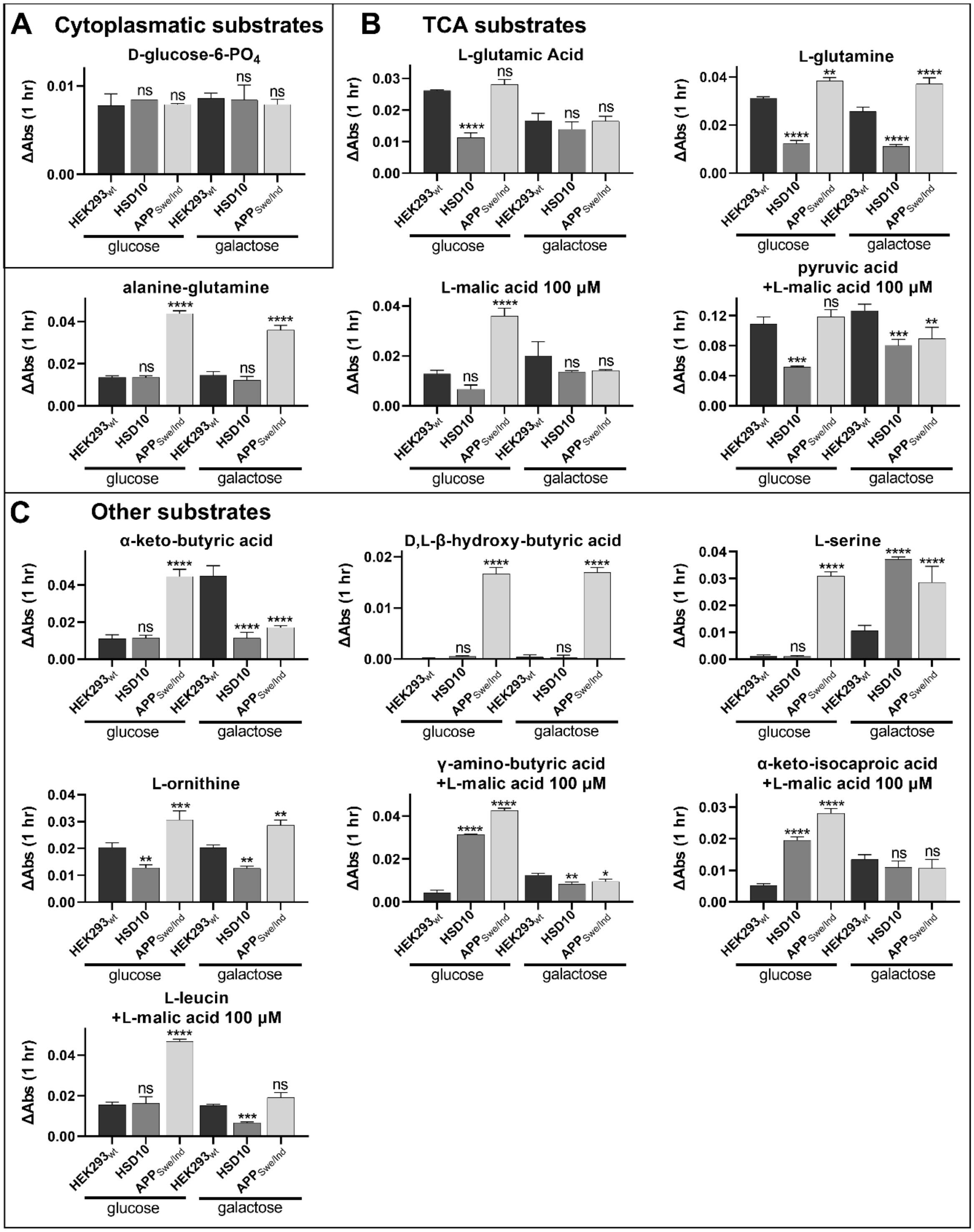


**Figure S15:** The mitochondrial electron flow changes (correspond to metabolic changes) in $HEK293_{wt}$, HSD10, and $APP_{Swe/Ind}$ cells measured by absorbance ($OD_{590}$) using MitoPlate assay. Conversion of cytosolic substrates (**A**), TCA cycle substrates (**B**), and other substrates (**C**) in $HEK293_{wt}$, HSD10, and $APP_{Swe/Ind}$ cells. Values are given as means ± SD from three independent cell culture preparations with one technical replicate (n=3). SD; standard deviation.

**Table S16:** ATP levels and cytotoxicity in $HEK293_{wt}$, and HSD10 cells performed 72 hr post-seeding into the galactose medium containing 7.6 nM Aβ42. Data were normalized between DMSO-treated (1%) and valinomycin-treated (100 µM) $HEK293_{wt}$ cells cultivated in galactose media. Values are given as means ± SD from three independent cell culture preparations with three technical replicates (n=9). SD; standard deviation.

| Cell line | Treatment | % of ATP quantity | % of Cytotoxicity |
|---|---|---|---|
| **$HEK293_{wt}$** | untreated | 100.00 ± 3.88 | 0.00 ± 0.63 |
| | 7.6 nM Aβ42 | 77.21 ± 4.23 | 13.87 ± 1.92 |
| **HSD10** | untreated | 74.94 ± 3.04 | 18.56 ± 2.82 |
| | 7.6 nM Aβ42 | 65.32 ± 3.71 | 25.28 ± 3.10 |

**Table S17:** ATP levels and cytotoxicity in HSD10 cells after HSD10-inhibitors treatment performed 72 hr post-seeding into the galactose medium. Data were normalized between DMSO-treated (1%) and valinomycin-treated (100 µM) $HEK293_{wt}$ cells cultivated in galactose media. Values are given as means ± SD from three independent cell culture preparations with four technical replicates (n=12). SD; standard deviation.

| Compound | | % of ATP quantity | % of Cytotoxicity |
|---|---|---|---|
| **DMSO** | 1% | 70.69 ± 3.68 | 18.08 ± 1.91 |
| **AG18051** | 94 nM | 85.94 ± 8.32 | 16.05 ± 2.56 |
| | 187 nM | 88.47 ± 7.05 | 14.00 ± 1.53 |
| | 0.38 µM | 89.20 ± 6.98 | 12.26 ± 1.87 |
| | 0.57 µM | 88.28 ± 5.23 | 9.24 ± 1.23 |
| | 0.75 µM | 83.09 ± 6.48 | 12.58 ± 1.33 |
| **34** | 2.13 µM | 85.47 ± 9.84 | 15.31 ± 1.03 |
| | 4.26 µM | 88.96 ± 8.08 | 14.63 ± 1.56 |
| | 8.52 µM | 88.42 ± 6.60 | 13.38 ± 1.07 |
| | 12.78 µM | 88.64 ± 7.53 | 10.26 ± 1.94 |
| | 17.04 µM | 84.94 ± 9.90 | 12.29 ± 3.04 |

**Table S18:** ATP levels and cytotoxicity in HSD10 cells performed 72 hr post-seeding into the Aβ42-containing galactose medium and HSD10-inhibitors treatment. Data were normalized between DMSO-treated (1%) and valinomycin-treated (100 µM) HEK293 cells cultivated in galactose media. Values are given as means ± SD from three independent cell culture preparations with three technical replicates (n=9). SD; standard deviation.

| Compound | % of ATP quantity | % of Cytotoxicity |
|---|---|---|
| **DMSO (1%)<br>Aβ42 (7.6 nM)** | 65.32 ± 3.71 | 25.28 ± 3.10 |
| **AG18051 (0.57 µM)<br>Aβ42 (7.6 nM)** | 65.77 ± 4.96 | 24.33 ± 3.72 |
| **34 (12.78 µM)<br>Aβ42 (7.6 nM)** | 71.50 ± 6.94 | 14.51 ± 3.71 |
